# Gaze and Pupil Patterns in ADHD and ASD during Naturalistic Viewing

**DOI:** 10.64898/2026.11.23.768639

**Authors:** Aimar Silvan, Vinay S. Raghavan, Jens Madsen, Lucas C. Parra

## Abstract

**Background:** Autism spectrum disorder (ASD) and attention-deficit/hyperactivity disorder (ADHD) frequently co-occur, and both are associated with atypical attention and arousal. Whether these associations arise from shared or distinct processes remains unclear because the conditions are rarely modeled jointly and attention and arousal are rarely measured simultaneously.

**Methods:** We analyzed eye-tracking data (gaze position and pupil size) collected during naturalistic video viewing from 2,421 youth (aged 5–22) in the Healthy Brain Network. ASD, ADHD-Inattentive, and ADHD-Hyperactive diagnostic statuses were modeled jointly alongside age and sex.

**Results:** Atypical gaze trajectories were uniquely associated with ASD. Basic oculomotor control and orienting after scene cuts and to faces were typical, but autistic participants maintained gaze consistently farther from typical viewing locations and spent less time looking at characters, faces, and mouths. Pupil fluctuations were atypical in both ASD and ADHD-Hyperactivity via distinct routes: reduced pupil synchrony in ASD was largely explained by gaze allocation, whereas in ADHD-Hyperactivity it was gaze-independent and accompanied by faster pupil fluctuations.

**Conclusions:** ASD and ADHD show dissociable signatures during natural viewing: ASD differs in what is looked at rather than when, whereas ADHD-Hyperactivity is characterized by faster-fluctuating arousal.

## 1. Introduction

Autism spectrum disorder (ASD) and attention-deficit/hyperactivity disorder (ADHD) frequently cooccur [1, 2], and both involve differences in attention [3, 4] and autonomic activity [5–7]. Neurocognitive theories explain such differences for each condition [8–10]. Yet, because the conditions are frequently studied separately without accounting for their co-occurrence, it remains difficult to determine which behavioral and physiological differences are specific to either diagnosis.

Prior eye-tracking and pupillometry studies in ASD and ADHD have often relied on constrained laboratory tasks using simple, abstract stimuli [11, 12]. While these tasks provide experimental control, they may not capture how individuals spontaneously respond to complex, continuously changing stimuli. Naturalistic video viewing offers an opportunity to investigate these responses with-out imposing explicit task demands [12, 13]. Eye tracking provides simultaneous measurements of gaze position and pupil size. Gaze position reflects overt visual attention, while pupil fluctuations are influenced by visual stimulation, autonomic activity, and cognitive processes such as attention and engagement [14, 15].

Atypical naturalistic gaze is linked to autistic traits [16–18], but its link to ADHD is less clear. Atypical gaze has been reported in ADHD, from impaired saccade inhibition in constrained tasks [19–21] to weaker top-down guidance during naturalistic viewing [22]. However, few of these studies model co-occurring autism, and those that do find that reduced social attention tracks autistic rather than ADHD traits [23, 24]. They also rarely consider age, although hyperactive-impulsive traits decline through childhood while inattentive traits persist [25, 26], and visual attention itself matures [27]. It remains unclear whether atypical gaze is independently associated with ADHD or confounded by ASD or age.

Naturalistic videos can elicit similar gaze patterns across viewers [28], allowing atypical gaze patterns to be quantified through inter-subject correlation (ISC), which measures the similarity of individual gaze responses to those of other viewers [28–30].

Autistic individuals look less at social content in dynamic scenes [31–33], but the underlying gaze behavior remains unclear. Findings on basic oculomotor control in ASD are conflicting [34, 35]. On static images, ASD is linked to higher center bias and reliance on visual saliency compared to adults without ASD [36]. Reduced social attention could stem from impaired orienting (slower first fixations to faces) [37, 38], or reduced maintenance (typical orienting followed by accumulating divergence) [39, 40]. Scene cuts in naturalistic videos provide an opportunity to examine how gaze behavior evolves following changes in visual content, including the timing of orienting and subsequent gaze allocation.

The locus coeruleus-norepinephrine (LC-NE) system contributes to the regulation of arousal and attention [14], with proposed alterations in both ASD and ADHD [41,42]. Adaptive gain theory suggests that phasic LC-NE activity supports focused engagement, while tonic activity promotes disengagement and exploration [14, 15]. Yet pupillometry findings are mixed: ASD studies report inconsistent pupil size and stimulus-evoked responses [43], while ADHD studies show smaller pupil sizes during attention-demanding tasks [44], potentially reflecting reduced arousal [6]. Direct comparisons report condition-specific profiles [45], with atypical arousal regulation in ASD but not ADHD [5].

These findings largely come from constrained tasks. Naturalistic stimuli can elicit synchronized pupil fluctuations across viewers (pupil ISC) [28, 29, 46]. Autistic children show reduced pupil synchrony during movie viewing, interpreted as altered arousal regulation [47, 48]. However, pupil differences could reflect varying visual input or differences in physiological dynamics not explained by gaze behavior. Beyond pupil synchrony and mean pupil size, the temporal structure of pupil fluctuations may provide additional information. In adults, the 1/ *f* exponent of resting pupil fluctuations predicts attentional orienting and self-rated hyperactivity-impulsivity, but not inattention, whereas mean pupil size, variance, and hippus power do not [49]. Whether the exponent differs across clinical diagnoses in youth or is independently associated with ADHD or ASD remains unknown.

To address these questions, we analyzed eye-tracking data collected during naturalistic video viewing in the Healthy Brain Network (HBN), a large pediatric cohort characterized by high prevalence of ASD, ADHD, and their co-occurrence [50, 51]. The scale and clinical diversity of this cohort allow us to examine unique associations with ASD and ADHD. Consistent with the Research Domain Criteria (RDoC) framework [52, 53], we jointly modeled ASD, ADHD-inattentive, and ADHD-hyperactive statuses alongside age and sex rather than using mutually exclusive groups. This dimensional approach isolates each diagnostic axis while holding overlapping traits and demographics constant, an approach previously shown to uncover condition-specific signatures in motor and speech behaviors [54].

We tested whether gaze atypicalities independently appear with ADHD, specified which gaze features (oculomotor control, spatial allocation, social attention, or orienting timing) are different in ASD, and characterized how pupillary synchrony and temporal dynamics diverge between conditions and relate to gaze.

## 2. Methods and Materials

### 2.1. Clinical Population & Diagnostic Procedure

This study analyzed eye-tracking data collected as part of the broader Healthy Brain Network effort [50, 51]. After strict quality control, 2,421 participants had usable data combined with clinician-confirmed diagnoses (Supplementary Section S1.1).

The cohort featured 1,374 individuals with clinician-confirmed ADHD, categorized into inattentive (668), hyperactive-impulsive (100), and combined (606) presentations. The sample included 280 individuals with ASD (>95% mild-to-moderate, primarily level 1), of whom 75.7% had co-occurring ADHD, at the upper end of community estimates [55, 56] and reflecting HBN’s clinical referral patterns. The study also included 246 typically developing (TD) participants (no DSM-5 diagnoses) and 626 individuals with other DSM-5 diagnoses, primarily anxiety and learning disorders, but without ADHD or ASD.

Differences in age remained generally small across the typically developing and diagnostic groups despite the broad developmental scope (5 to 22 years old, *M* = 10.35, *SD* = 3.45, Table 1). The male-to-female ratio was 1.55:1 (1,472:949).

**Table 1.**
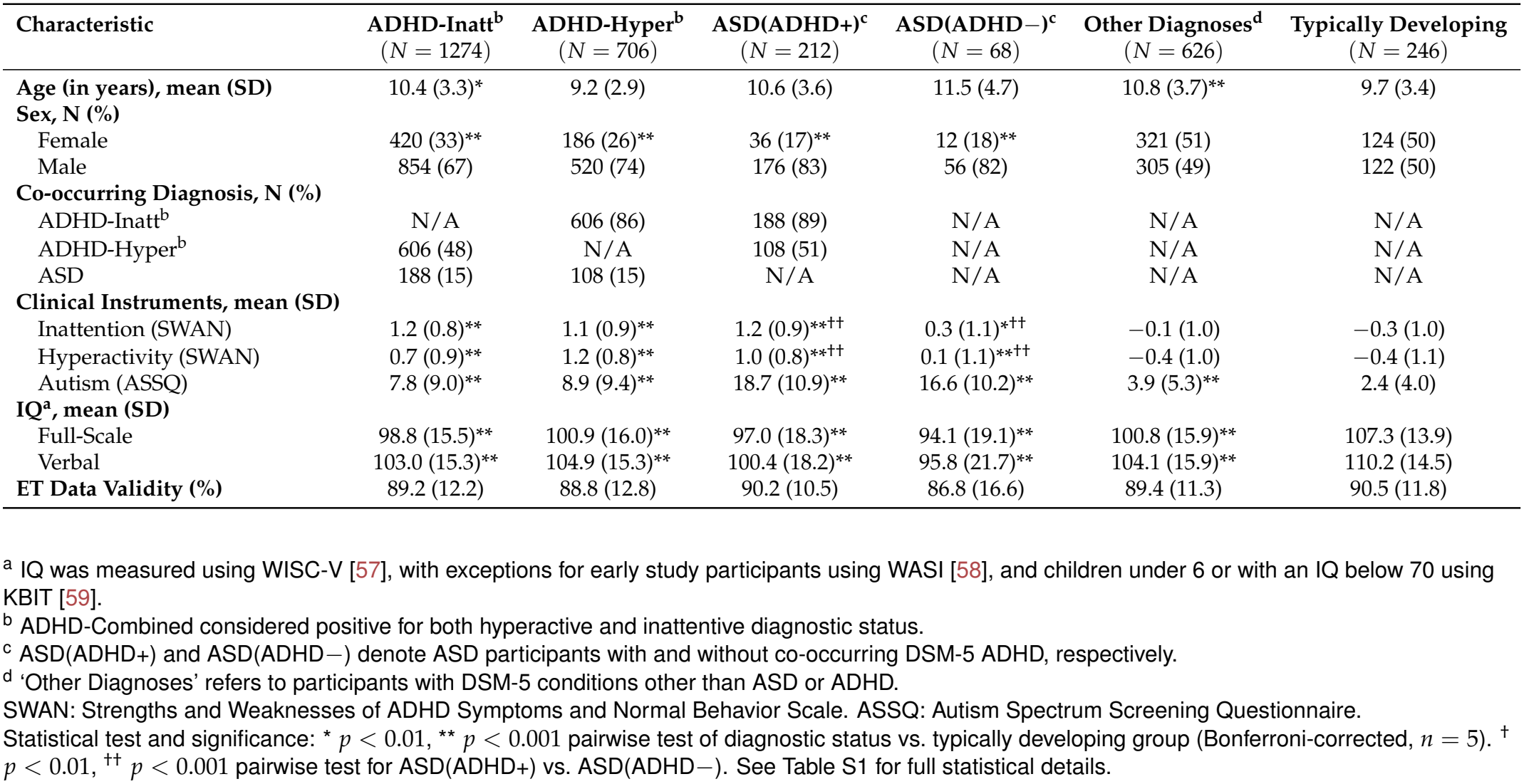
Demographic and clinical characteristics.

| Characteristic | ADHD-Inatt<sup>b</sup><br>(*N* = 1274) | ADHD-Hyper<sup>b</sup><br>(*N* = 706) | ASD(ADHD+)<sup>c</sup><br>(*N* = 212) | ASD(ADHD−)<sup>c</sup><br>(*N* = 68) | Other Diagnoses<sup>d</sup><br>(*N* = 626) | Typically Developing<br>(*N* = 246) |
|---|---|---|---|---|---|---|
| **Age (in years), mean (SD)** | 10.4 (3.3)<sup>*</sup> | 9.2 (2.9) | 10.6 (3.6) | 11.5 (4.7) | 10.8 (3.7)<sup>**</sup> | 9.7 (3.4) |
| **Sex, N (%)** |  |  |  |  |  |  |
| Female | 420 (33)<sup>**</sup> | 186 (26)<sup>**</sup> | 36 (17)<sup>**</sup> | 12 (18)<sup>**</sup> | 321 (51) | 124 (50) |
| Male | 854 (67) | 520 (74) | 176 (83) | 56 (82) | 305 (49) | 122 (50) |
| **Co-occurring Diagnosis, N (%)** |  |  |  |  |  |  |
| ADHD-Inatt<sup>b</sup> | N/A | 606 (86) | 188 (89) | N/A | N/A | N/A |
| ADHD-Hyper<sup>b</sup> | 606 (48) | N/A | 108 (51) | N/A | N/A | N/A |
| ASD | 188 (15) | 108 (15) | N/A | N/A | N/A | N/A |
| **Clinical Instruments, mean (SD)** |  |  |  |  |  |  |
| Inattention (SWAN) | 1.2 (0.8)<sup>**</sup> | 1.1 (0.9)<sup>**</sup> | 1.2 (0.9)<sup>**††</sup> | 0.3 (1.1)<sup>*††</sup> | −0.1 (1.0) | −0.3 (1.0) |
| Hyperactivity (SWAN) | 0.7 (0.9)<sup>**</sup> | 1.2 (0.8)<sup>**</sup> | 1.0 (0.8)<sup>**††</sup> | 0.1 (1.1)<sup>**††</sup> | −0.4 (1.0) | −0.4 (1.1) |
| Autism (ASSQ) | 7.8 (9.0)<sup>**</sup> | 8.9 (9.4)<sup>**</sup> | 18.7 (10.9)<sup>**</sup> | 16.6 (10.2)<sup>**</sup> | 3.9 (5.3)<sup>**</sup> | 2.4 (4.0) |
| **IQ<sup>a</sup>, mean (SD)** |  |  |  |  |  |  |
| Full-Scale | 98.8 (15.5)<sup>**</sup> | 100.9 (16.0)<sup>**</sup> | 97.0 (18.3)<sup>**</sup> | 94.1 (19.1)<sup>**</sup> | 100.8 (15.9)<sup>**</sup> | 107.3 (13.9) |
| Verbal | 103.0 (15.3)<sup>**</sup> | 104.9 (15.3)<sup>**</sup> | 100.4 (18.2)<sup>**</sup> | 95.8 (21.7)<sup>**</sup> | 104.1 (15.9)<sup>**</sup> | 110.2 (14.5) |
| **ET Data Validity (%)** | 89.2 (12.2) | 88.8 (12.8) | 90.2 (10.5) | 86.8 (16.6) | 89.4 (11.3) | 90.5 (11.8) |

<sup>a</sup> IQ was measured using WISC-V [57], with exceptions for early study participants using WASI [58], and children under 6 or with an IQ below 70 using KBIT [59].
<sup>b</sup> ADHD-Combined considered positive for both hyperactive and inattentive diagnostic status.
<sup>c</sup> ASD(ADHD+) and ASD(ADHD−) denote ASD participants with and without co-occurring DSM-5 ADHD, respectively.
<sup>d</sup> ‘Other Diagnoses’ refers to participants with DSM-5 conditions other than ASD or ADHD.
SWAN: Strengths and Weaknesses of ADHD Symptoms and Normal Behavior Scale. ASSQ: Autism Spectrum Screening Questionnaire.
Statistical test and significance: <sup>\*</sup> *p* < 0.01, <sup>\*\*</sup> *p* < 0.001 pairwise test of diagnostic status vs. typically developing group (Bonferroni-corrected, *n* = 5). <sup>†</sup> *p* < 0.01, <sup>††</sup> *p* < 0.001 pairwise test for ASD(ADHD+) vs. ASD(ADHD−). See Table S1 for full statistical details.

In HBN, licensed clinicians establish consensus DSM-5 diagnoses primarily via the semi-structured Kiddie Schedule for Affective Disorders and Schizophrenia (K-SADS) [60] with youth and guardians, integrating clinical testing observations, standardized questionnaires, and developmental and academic history.

ASD assessment primarily uses the K-SADS autism module and specific questionnaires (SRS-2, SCQ, ASSQ), with only ~ 20 participants ADOS-confirmed. To ensure diagnostic certainty, individuals recommended for additional testing were excluded from analysis. Detailed protocols are in the original HBN publications [50, 51].

The New York-based community sample was racially diverse: 44.89% White/Caucasian, 9.77% Black/African American, 7.44% Hispanic, 2.21% Asian, 0.69% Indian, 1.68% other races, 13.25% multiracial, and 20.07% unknown or unspecified.

Only ~ 1% of participants were under active stimulants on testing day.

### 2.2. Stimuli, Data Acquisition, and Pre-processing

**Stimuli**. Participants passively viewed four ageappropriate video clips: “Despicable Me” (DM, 2:50 min), “Fun with Fractals” (FF, 2:39 min), “Diary of a Wimpy Kid” (WK, 1:57 min), and “The Present” (TP, 4:18 min; stimulus links in Supplementary Section S1.1).

#### Acquisition

Eye position and pupil size were recorded in a sound-shielded room at ~ 700 mm distance using a chin rest and an infrared eye tracker (iView-X Red-m, SMI GmbH; 30, 60, or 120 Hz sampling rate; 0.1*°* resolution; 0.5° accuracy). Prior to each video, calibration used a 5-point grid and was repeated until maximum point error fell below 2° or average error fell below 1° [51].

#### Eye-tracking pre-processing and event detection

Only maximum-confidence gaze samples were retained; missing samples were linearly interpolated, and time series were resampled to 30 Hz. Video views with <65% valid pre-interpolation data, >65% off-stimulus gaze, or duration deviations >0.5 s were excluded (Supplementary Section S1.1–S1.2). Using the eye with higher validity, blinks were identified with default eye-tracker settings, and fixations and saccades were detected from 3D gaze vectors using an Identification by Dispersion-Threshold (I-DT) algorithm [22, 61]. From these events, we computed per-stimulus mean fixation duration, mean saccade amplitude, blink rate, and coefficients of variation (CV) for fixation duration and saccade amplitude (Supplementary Section S1.3).

#### Pupil pre-processing

To isolate cognitive and autonomic pupil fluctuations from mechanical and visual artifacts, we regressed out blink offsets, saccade onsets, saccade amplitude, log-luminance, and luminance change per participant using sequential banded ridge Temporal Response Functions (TRFs; − 4 to 6 s lag window) via the mTRF-Toolbox [46, 62, 63]. Residual time series served as the cleaned pupil signal for Pupil-ISC and spectral analyses (full TRF formulation and cross-validation in Supplementary Section S1.4).

### 2.3. Inter-Subject Correlation (ISC)

To quantify typical gaze and pupil trajectories, Inter-Subject Correlation (ISC) was calculated for the continuous time-series. Reference patterns were constructed for each stimulus by calculating the median gaze position or pupil size across the typically developing (TD) cohort at each time point (using a leave-one-out approach for TD participants).

Gaze-ISC was defined as the average Pearson correlation between a participant’s continuous gaze coordinates (*x* and *y*) and the reference trajectories. Pupil-ISC was similarly calculated using the artifact-cleaned pupil signal. All ISC values were Fisher-*z* transformed prior to statistical analysis to approximate normality.

### 2.4. Visual feature extraction

High- and low-level visual features were extracted from videos to measure their impact on viewing patterns.

Regions of interest (ROIs) including faces, characters (human, puppy), mouth, and eyes were automatically segmented using SAM3 [64]. Text was detected with easyocr [65], and unsegmented areas formed the back-ground. “Despicable Me” and “The Present” featured animated characters; “Diary of a Wimpy Kid” contained live-action actors; and “Fun with Fractals” contained text only.

#### Visual Saliency

Per-frame pixel-level saliency maps were generated using the Itti-Koch model [66], combining intensity, color, and orientation into normalized probability distributions.

Luminance was calculated as mean lightness in CIE L*a*b* space. Scene cuts were identified via temporal contrast peaks [67] and manually verified.

### 2.5. Gaze allocation measures

#### Latency to fixation after scene cuts

Cut-to-fixation latency was measured for first fixations beginning within 1 s post-cut. Latencies were log-transformed and averaged per participant and stimulus.

#### Region-of-interest preference

ROI masks were dilated ~1° to account for eye-tracking inaccuracy. Preference was defined as 100*×* (frames gaze on ROI / frames ROI present). First-fixation latency to faces post-cut was evaluated for cuts with an initial face ROI. Latencies were log-transformed and averaged per participant and stimulus.

Center bias was the percentage of valid gaze samples within 5° of the stimulus center. Following Wang et al. [36], we also measured an object-free subset: frames where no ROI intruded into a central 2° disc.

#### Saliency at fixation

We calculated the within-frame percentile rank of the saliency map at each fixation mid-point (chance = 0.5). Maps were blurred (~ 1° Gaussian) for eye-tracking inaccuracy.

#### Gaze distance around scene cuts

We measured the Euclidean distance (° visual angle) between a participant’s gaze and the median gaze of TD controls (leave-one-out).

Distances were measured− 1 to 3 s relative to scene cuts and averaged per participant and stimulus.

### 2.6. 1/*f* exponent of pupil fluctuations

Frequency content was quantified from the artifact- and luminance-cleaned pupil time series. Spectral power was estimated over a 0.02–2.0 Hz band [49] and fitted in log-log space as:

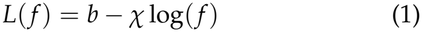

where *b* is the broadband offset and *χ* is the exponent (spectral drop-off slope) [68].

### 2.7 Analytical Approach & Statistical Modeling

To evaluate relationships between target measurements and clinical diagnoses while controlling for confounders, we used Linear Mixed-Effects (LME) models to account for stimulus-level variance and baseline individual differences across repeated observations:

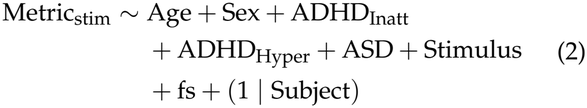

Predictors included continuous age, sex (Female=1, Male=0), clinician-confirmed diagnoses (ASD, ADHD-Inattentive, ADHD-Hyperactive; ADHD-Combined modeled as positive for both), and sampling rate (fs) as a categorical covariate for acquisition differences.

In this framework, each participant is a single subject with clinical coding. For example, a 9-year-old female with negative ASD and positive ADHD-Combined is coded as: 9 (Age), 1 (Sex), 0 (ASD), 1 (ADHD-Inatt), 1 (ADHD-Hyper).

Prior to model fitting, dependent variables and continuous predictors were z-scored, while binary diagnostic variables, sex, and sampling rate (fs) remained dummy-coded. Models were fitted using restricted maximum likelihood (REML) in MATLAB, with low multicollinearity confirmed (all VIFs < 1.32).

Because outcomes and continuous predictors were z-scored, covariate-adjusted *β* coefficients represent standardized mean differences (analogous to Cohen’s *d*) for binary predictors and standardized regression slopes (SD change per SD) for continuous predictors. Full model statistical tables are in Supplementary Section S3.

To test the robustness of our conclusions, we conducted several post-hoc sensitivity analyses, each using the same linear mixed-effects framework and covariates as the main model (Supplementary Section S2).

Statistical significance was set at *α* = 0.01, and standardized effect sizes (*β*) are reported in the text for significant associations. Bonferroni correction was applied within multi-outcome families: basic oculomotor means (*n* = 4), oculomotor coefficients of variation (*n* = 2), content ROIs (*n* = 6), and added-covariate Pupil-ISC models (*n* = 3).

To quantify diagnostic associations, we used logistic regressions controlling for age, sex, and the remaining diagnostic variables. Age and sex aORs came from each diagnosis’s respective model; for each diagnostic pair, we averaged the two log odds ratios and standard errors to derive a single symmetric aOR and applied Bonferroni correction across the three tests.

## 3. Results

### 3.1. Diagnostic Co-occurrence and Demographic Associations

Across the 2,421 participants who completed the naturalistic viewing paradigm (Fig. 1a; Table 1, Fig. S1), diagnostic statuses showed substantial overlap and demographic skew (Fig. 1b; Table S2). ADHD-Inattentive status was associated with higher odds of co-occurring ASD (aOR = 1.62), whereas ADHD-Hyperactive status was not (aOR = 1.22, n.s.); the two ADHD presentations were strongly linked (aOR = 10.70). Odds of an ADHD-Hyperactive diagnosis decreased with age (aOR = 0.81/year), while odds of ADHD-Inattentive and ASD diagnoses increased (aOR = 1.08 and 1.06/year). Females had lower odds across all diagnoses, particularly ASD (aOR = 0.30). All subsequent analyses therefore jointly modeled all three diagnostic statuses alongside age and sex.

**Figure 1:**
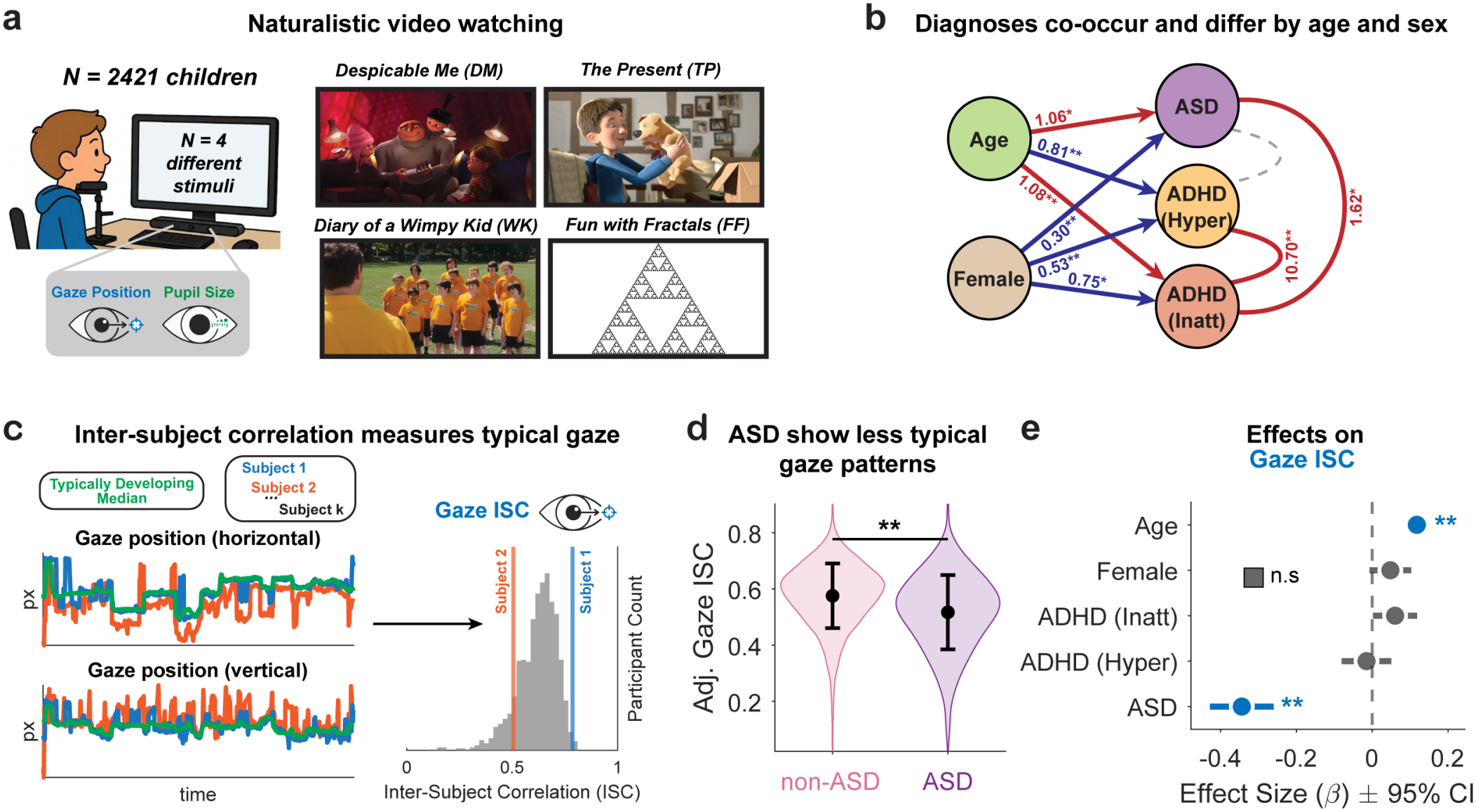
Inter-subject correlation captures atypical viewing patterns in ASD but not ADHD. (a) *N* = 2421 children and young adults, aged 5–22, watched four videos while gaze position and pupil size were recorded. See Table 1 for specific demographic, clinical, and trait characterization. (b) Arrows point from age and sex to each diagnostic status and show the adjusted odds ratio (aOR) for that status. Lines between diagnoses show symmetric aORs, adjusted for the third diagnosis, age and sex. Red, higher odds; blue, lower odds; dashed grey, not significant. (* *p* < 0.01, ** *p* < 0.001, Bonferroni-corrected; see Table S2). (c) Inter-subject correlation (ISC) is a proxy for typical viewing patterns. Horizontal and vertical gaze position time series for individual participants (e.g., Subject 1, Subject 2) correlate with the median trajectory of the Typically Developing references (green line). The distribution of ISC values reflect how typical each participant’s viewing patterns are (Gaze-ISC). (d) Partial dependence plot demonstrating the effect of a positive Autism Spectrum Disorder (ASD) clinical diagnosis on Gaze-ISC. Bars = mean ± std, ** *p* < 0.001. (e) Forest plot summarizing the standardized effect sizes (*β*) ± 95% CI for Gaze-ISC. Significant associations are colored (** *p* < 0.001), gray if non-significant.

### 3.2. Atypical gaze patterns are unique to ASD; association with ADHD is due to age effects

We first asked whether atypical naturalistic gaze relates to ASD, ADHD, or both. An ADHD-specific effect predicts gaze disruption independent of other diagnoses, but apparent effects might reflect confounding by co-occurring ASD or by developmental stage, since ADHD presentations differ and change with age (Fig. 1b).

In the full model of Gaze-ISC (Fig. 1c), ASD was independently associated with lower Gaze-ISC (*β* = 0.344; Fig. 1d, e), and older age predicted higher Gaze-ISC (*β* = − 0.118; Fig. 1e). Controlling for ASD, age, and sex, neither ADHD presentation significantly predicted Gaze-ISC (Fig. 1e).

Refitting models without covariates showed how spurious ADHD effects emerge (Fig. S2). Omitting age and sex yielded small, opposite-signed ADHD effects: Hyperactive predicted lower Gaze-ISC (*β* = − 0.090), and Inattentive predicted higher Gaze-ISC (*β* = 0.085). Mirroring the age differences across presentations (Fig. 1b):

Hyperactivity is more prevalent in younger and inattention in older participants, and Gaze-ISC increases with age. Omitting ASD yielded no significant ADHD effects. Thus, apparent ADHD effects on gaze typicality arose from age confounding rather than ADHD itself, supporting use of the full joint model throughout.

### 3.3. ASD gaze differs in what is looked at, not in oculo-motor control or orienting latency

We next tested whether reduced Gaze-ISC in ASD reflects differences in basic oculomotor control, spatial strategy, pixel-level saliency, social content selection, or orienting timing around scene cuts.

Basic oculomotor measures (fixation count/duration, blink rate, saccade amplitude; Fig. 2a) and exogenous orienting latency after scene cuts (Fig. 2b) showed no diagnostic associations, including for within-participant variability (Fig. S3). Older age predicted higher blink rates (*β* = 0.172), smaller saccades (*β* = − 0.088), and faster orienting (*β* =− 0.04). Thus, reduced gaze syn-chrony in ASD does not seem to stem from basic oculo-motor control or exogenous orienting.

**Figure 2:**
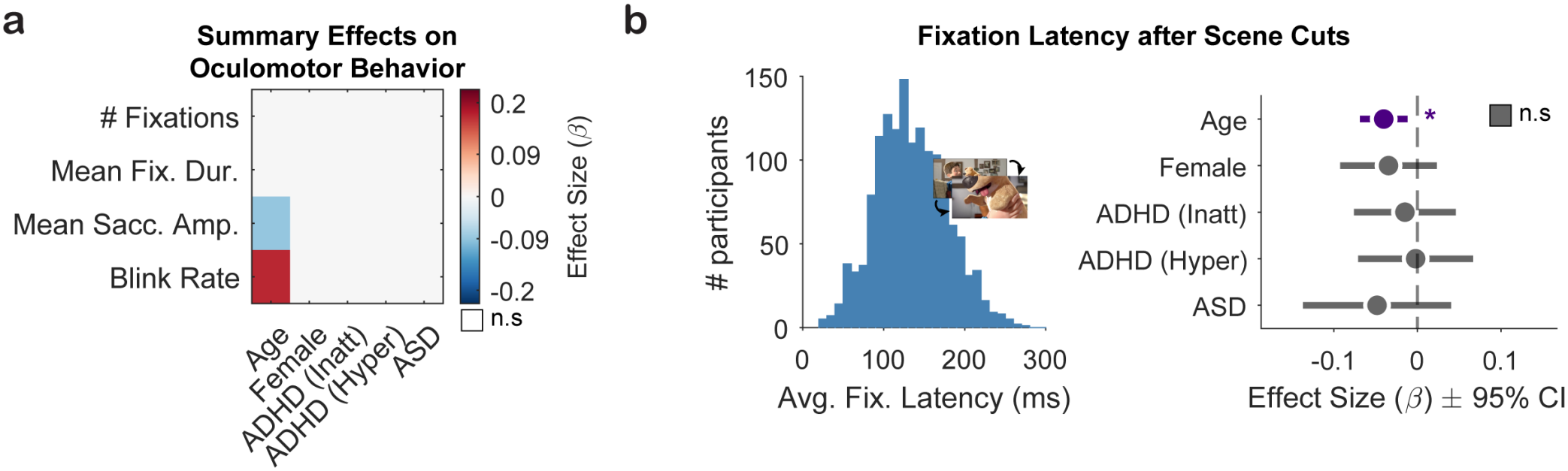
Basic oculomotor behaviors remain typical in ADHD and ASD. (a) Heatmap depicting the standardized effect sizes (*β*) for basic oculomotor behaviors. Colored cells indicate significant main effects (*p* < 0.01, Bonferroni-corrected), white otherwise. (b) Left: Average fixation latency after scene cuts across participants when watching “The Present”. Right: forest plot showing the adjusted effects (*β* ± 95% CI) predicting fixation latency after scene cuts. Significant features are shown in purple (* *p* < 0.01). Summary statistics for panels a–b are available in Supplementary Section S3.

We next examined spatial gaze allocation and content features (Fig. 3a). Contrary to expectations, ASD was associated with reduced center bias (*β* = − 0.128; Fig. 3b), a finding that persisted even in scenes lacking central objects (*β* = − 0.124; Fig. S4). Older age was associated with greater center bias (*β* = 0.073). Fixated pixel-level saliency [66] was not associated with any diagnosis (Fig. 3c).

**Figure 3:**
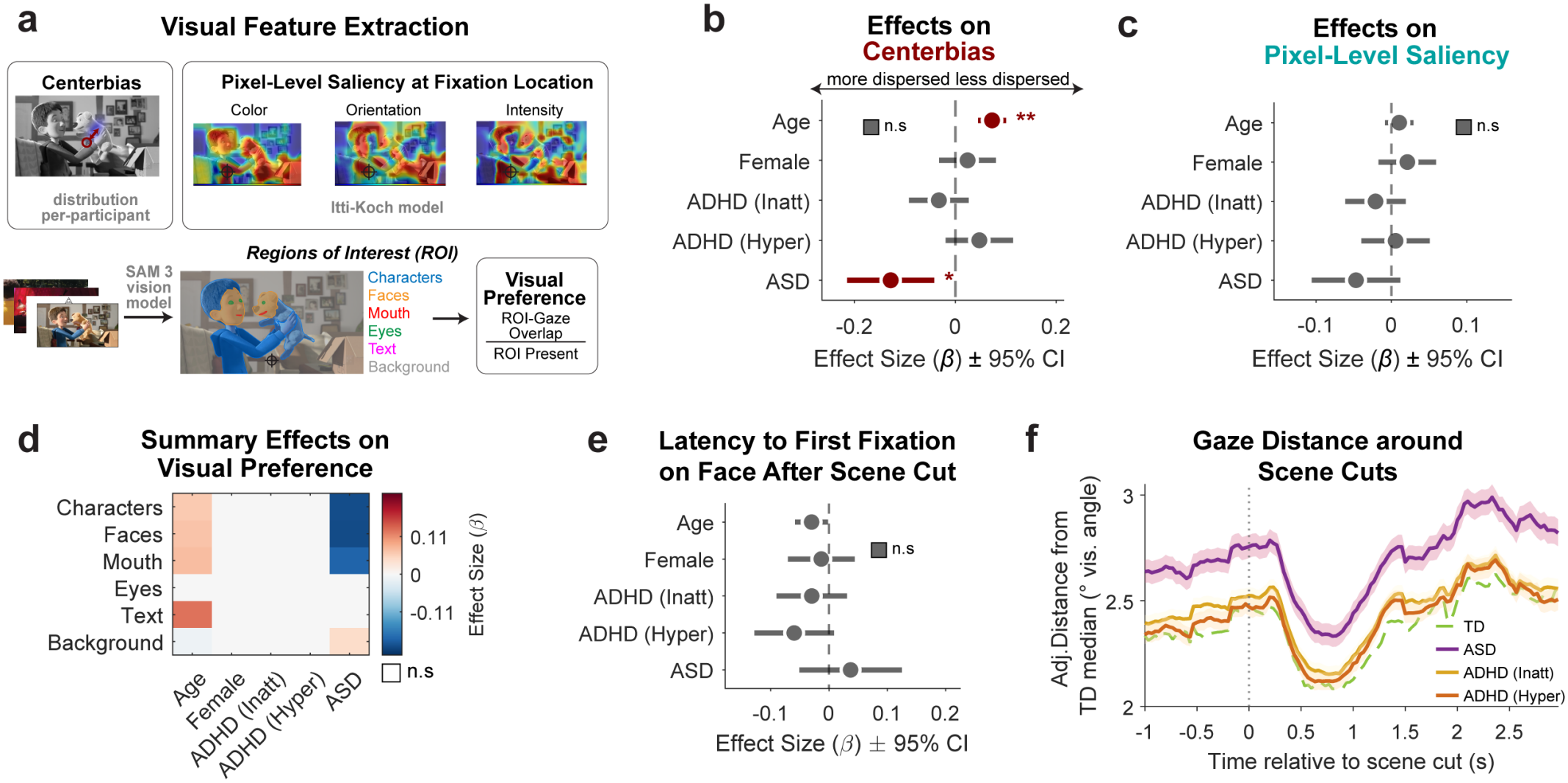
ASD gaze differs in what is looked at, not in oculomotor control, orienting, or timing after scene cuts. (a) Features to measure spatial gaze allocation. Center bias was calculated as the percentage of valid gaze samples falling within 5° of the stimulus center. Pixel-level saliency was computed using the Itti-Koch model [66]. Regions of interest (ROIs) were segmented using the SAM3 vision model [64]. (b) Forest plot depicting effects on center bias. (c) Forest plot of effects for average pixel-level saliency at fixation. (d) Heatmap summarizing standardized effect sizes *β* for visual preference, defined as the percentage of time gaze fell within an ROI when present. Colored cells indicate significant main effects (*p* < 0.01, Bonferroni-corrected). (e) Forest plot of effects on the latency to the first fixation on a face after a scene cut. (f) Distance of gaze from the median gaze of typically developing participants, in degrees of visual angle, resolved by time from scene cuts (dotted line, *t* = 0). For TD participants, the median was computed leaving the participant out. Lines show mean± SE of covariate-adjusted distance predictions. Panels b, c, and e depict standardized effect sizes (*β* ±95% CI). Significant associations are colored (* *p* < 0.01, ** *p* < 0.001), gray if non-significant. Summary statistics for panels b–e are available in Supplementary Section S3.

Using content-defined ROIs, ASD was associated with less total fixation time on characters (*β* =− 0.210), faces (*β* = −0.212), and mouths (*β* =− 0.188; Fig. 3d). However, initial face fixation latency after scene cuts did not differ (Fig. 3e). Older participants showed stronger ROI preferences, particularly for text (*β* = 0.129).

Tracking gaze distance from the TD median following scene cuts (Fig. 3f) revealed that while all groups converged within~ 1 second before dispersing, ASD gaze remained consistently farther from typical locations throughout. Thus, ASD divergence is not time-locked to scene cuts, and orienting seems typical.

Thus, ASD gaze differed in what was looked at rather than in when, while neither ADHD presentation showed associations with any oculomotor, spatial, saliency, or ROI measure.

### 3.4. Pupil fluctuations are atypical in both ASD and ADHD-Hyperactive, but only hyperactivity shows faster pupil fluctuations

Lastly, we asked whether pupil size fluctuations differ during naturalistic viewing. We measured this at three levels: stimulus-locked fluctuations (Pupil-ISC), overall size (mean pupil size), and the fluctuation speed (the 1/ *f* exponent of the pupil power spectrum).

Fitted nuisance TRFs (Fig. 4a) confirmed expected post-blink, post-saccade, and pupillary light reflex responses, with each regressor improving cross-validated prediction (Fig. S5). Using the cleaned pupil signal, both ASD (*β* = −0.141) and ADHD-Hyperactive status (*β* = −0.076) independently predicted lower Pupil-ISC (Fig. 4b), as did older age (*β* = 0.052).

**Figure 4:**
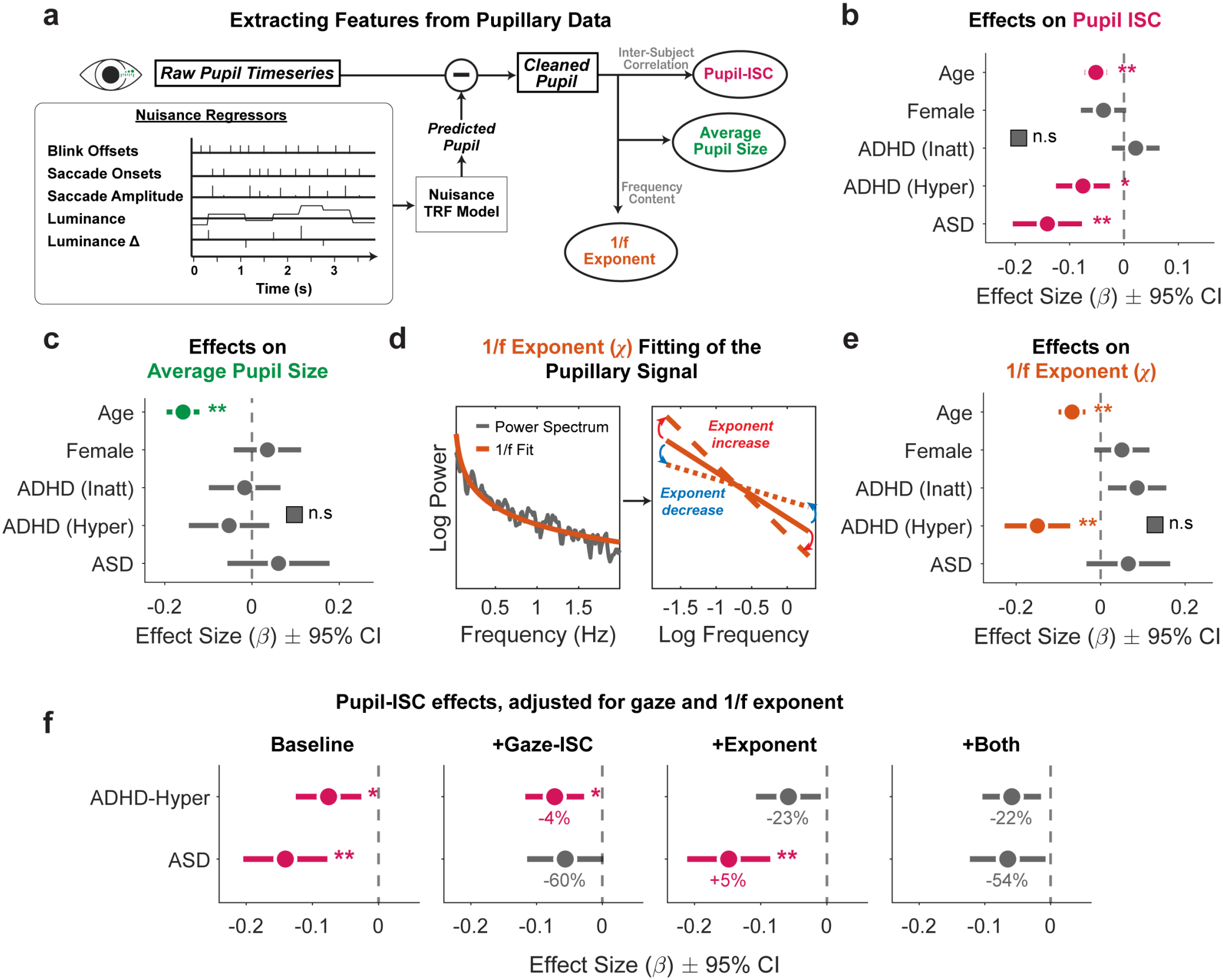
Parameterization and clinical associations of the pupillary dynamics. (a) Pupil size time series are residualized against a nuisance TRF model that incorporates oculomotor and low-level visual covariates to isolate a cleaned autonomic pupil signal, which is then used to parametrize typical pupil patterns (Pupil-ISC), mean pupil size, and the 1/ *f* exponent. (b) Forest plot depicting effects on Pupil-ISC, calculated by comparing each participant’s cleaned pupil signal to the median of the TD population. (c) Forest plot of multivariate effects on the average pupil size. (d) The power spectral density (PSD, gray line) of the cleaned pupil is fit to extract the 1/ *f* exponent (*χ*) in the 0.02–2 Hz range. Linear fit in log-log space (orange). A lower exponent *χ* measures more relative higher frequency components. (e) Forest plot of effect sizes on the 1/ *f* exponent. (f) Attenuation of Pupil-ISC effects by gaze and by the 1/ *f* exponent. Each panel shows the ADHD-Hyperactive and ASD effects on Pupil-ISC from the baseline model (as in b) and from models adding Gaze-ISC, the 1/ *f* exponent, or both as covariates. Percentages give the change in *β* relative to baseline; negative values indicate attenuation toward zero. values for the three added-covariate models are Bonferroni-corrected. Panels b, c, e, and f depict standardized effect sizes (*β* ± 95% CI). Significant associations are colored (* *p* < 0.01, ** *p* < 0.001), gray if non-significant. Summary statistics for b, c, and e are available in Supplementary Section S3.

Mean pupil size was unaffected by diagnoses and only decreased with age (*β* = − 0.158, Fig. 4c).

In spectral analysis of the cleaned pupil signal (Fig. 4d), ADHD-Hyperactive status (*β* = −0.150; Fig. 4e) and older age (*β* =− 0.068) predicted a lower 1/ *f* exponent *χ* (flatter spectrum, faster fluctuations), whereas ASD and ADHD-Inattentive status showed no association. Data validity (Table 1), and blink rates (Fig. 2a) did not differ for hyperactive participants, ruling out data loss or oculomotor artifacts.

Because both ASD and ADHD-Hyperactive status were associated with lower Pupil-ISC, we tested whether these effects arise through the same route. Adding Gaze-ISC as a covariate reduced the ASD effect on Pupil-ISC by 60% (*β* =− 0.141 to −0.056, n.s.) while leaving the hyperactivity effect unchanged (*β* =− 0.076 to− 0.073; Fig. 4f). Conversely, controlling for the 1/ *f* exponent reduced the yperactivity effect by 23% (to −0.058, n.s.) without affecting the ASD effect. Including both covariates reduced the ASD effect by 54% and yielded a hyperactivity effect similar to exponent-only control (*β* =− 0.059, n.s.). Thus, reduced pupil synchrony seems to arise from distinct mechanisms: from spatial gaze allocation in ASD, and partly from faster pupil fluctuations in hyperactivity.

## 4. Discussion

We investigated gaze behavior and pupil fluctuations in ASD and ADHD during natural viewing to identify mechanisms underlying attention and arousal. Jointly modeling ASD, ADHD presentations, age, and sex during naturalistic viewing revealed three findings. First, atypical gaze was unique to ASD, whereas apparent ADHD effects reflected age differences. Second, ASD gaze differed in content selection (less time on characters, faces, and mouths; greater distance from typical gaze) rather than oculomotor control or orienting timing. Third, both ASD and ADHD-Hyperactivity showed reduced pupil synchrony, but via distinct routes: viewed content in ASD versus gaze-independent, faster pupil fluctuations in hyperactivity.

Reduced gaze synchrony in ASD replicates prior ISC work [16, 17]. Gaze synchrony increased with age, reflecting normal visual development [69–71]. This age effect explains previous ADHD gaze reports comparing ADHD to controls [22]. Such comparisons are sensitive to age differences between presentations: hyperactive participants are typically younger and inattentive ones older, so ignoring age or pooling presentations can create or mask age-driven effects, as shown by our reduced models. While ADHD saccade inhibition deficits vary with age in task-constrained settings [72, 73], free viewing requires no inhibition, and basic saccade generation remains intact in ADHD [74, 75]. Thus, while task-driven gaze control differences may exist in ADHD, free-viewing atypical gaze is specific to ASD.

If gaze atypicality is specific to ASD, where does it arise? We found no ASD differences in basic oculomotor control or exogenous orienting after scene cuts. Prior free-viewing evidence is mixed: basic oculomotor function was typical in two large pediatric samples watching animated videos [34, 76], whereas reduced saccade amplitude and duration occurred in participants aged 6–30 [77], consistent with meta-analytic evidence of saccade dysmetria [35]. Higher blink rates have also been reported in younger autistic children [78, 79]. Given the similar age range in Bast et al. [77], age alone does not explain these discrepancies. Co-occurring ADHD is a likely candidate; most of our autistic participants had ADHD, and supplementary analyses revealed more variable saccade amplitudes only in those without ADHD, suggesting oculomotor profiles in autism depend on co-occurring conditions [80]. Typical orienting post-cut aligns with preserved simple target engagement and disengagement in ASD [35], indicating gaze divergence originates above low-level oculomotor control.

At the levels of spatial strategy and saliency, findings diverge from previous static image work [36]. ASD was linked to reduced, rather than increased, center bias, a spatial strategy that persisted even in scenes without central objects. Pixel-level saliency did not differ across groups, contrasting static image findings [36] but aligning with video research showing weaker low-level saliency prediction of autistic gaze [76].

Regarding social content, typical face latency and post-scene-cut gaze time courses, combined with a constant offset from typical viewing, indicate lower social interest rather than impaired orienting. Intact face orienting matches prior ASD reports [39, 40, 81]. Although slower orienting is reported elsewhere [37, 38], face prioritization depends on competing dynamic content [82, 83], which changes continuously during videos. We observed no intra-scene divergence accumulation within our multi-second analysis window around scene cuts, though slower video-wide drifts remain possible [39, 84].

Contrary to interpretations of reduced autistic pupil synchrony as idiosyncratic arousal regulation [47, 48], our results suggest it mainly reflects viewed content. In hyperactivity, reduced pupil synchrony was gazeindependent and partly driven by faster pupil fluctuations, supporting accounts of unstable arousal in ADHD [10, 85], EEG evidence of unstable regulation [86, 87], and hyperactive phasic LC responses [88]. Bast et al. [5] found atypical regulation in ASD instead of ADHD, likely due to task-evoked designs and pooling ADHD presentations, which masked hyperactivity effects in our data (Supplementary Section S2).

The 1/ *f* exponent captures this difference: resting pupil exponent reductions correlate with hyperactivityimpulsivity but not inattention in adults [49], a finding we extend to clinical diagnoses in youth and free viewing. Although EEG 1/ *f* findings in ADHD vary by age [89–91], age-related exponent flattening matches developmental EEG spectrum changes [89, 92]. Distinct from cortical excitation/inhibition in EEG, the pupillary 1/ *f* exponent reflects the balance of slow-tonic and fast-phasic fluctuations, shifted toward faster speeds in hyperactivity.

Age had consistent effects across all measures. Older participants showed higher gaze synchrony, greater center bias, and increased ROI preference, consistent with visual receptive field maturation [93]. In contrast to gaze, pupil synchrony decreased with age, possibly reflecting altered adolescent affective reactivity [94] or fluctuation speed.

Broadly, our results demonstrate how diagnostic overlap can mislead. ADHD presentations differ systematically in age [95], presentation shifts occur during development [25], case-control effects vary with age [96], and unmeasured traits can confound comparisons [23, 97]. Jointly modeling diagnoses with age and sex resolves these confounds across the entire sample. Results remained robust across six alternative analysis approaches (Supplementary Section S2), though pooling ADHD presentations or using parent ratings eliminated hyperactivity’s pupil synchrony effects, illustrating how pooling obscures presentation-specific findings. Co-occurring ASD and ADHD may also interact non-additively [48,98]: gaze atypicality was more pronounced in autistic participants without inattentive ADHD.

Study limitations should be noted. First, the cross-sectional nature impedes describing developmental trajectories. Second, caution is needed when generalizing these ASD findings to the broader autism spectrum. Participants had relatively high IQ (Table 1), so findings may not extend to those with intellectual disability or higher support needs. Additionally, 76% of the ASD sample had co-occurring ADHD, exceeding pooled estimates [2] reflecting HBN referral patterns. Although our joint model separates both diagnoses statistically, the findings apply most directly to this co-occurring profile. Third, TD participants had higher IQ. While IQ did not moderate ASD gaze differences in a meta-analysis [33], it may in select samples [99]. Here, higher IQ correlated with increased gaze and pupil synchrony, and including it slightly attenuated ASD effects without altering conclusions (Supplementary Section S2). Fourth, variable eye-tracker sampling rates may constrain saccade precision, though sampling rate was included as a covariate in all models. Fifth, mean pupil size was not referenced to a baseline and reflects viewing distance and session drift rather than tonic arousal, so we draw no inferences from its null result. Finally, effect sizes were small, derived strictly during passive viewing, and represent group-level markers rather than diagnostic instruments.

While autism and ADHD share surface features, a few minutes of naturalistic viewing separate them into distinct channels: gaze (reflecting attention allocation) and pupil dynamics (reflecting arousal fluctuations). Autism uniquely alters gaze patterns, whereas hyperactivity drives faster pupil fluctuations. These specific markers emerge only when co-occurring diagnoses, age, and sex are modeled jointly. Jointly modeling overlapping conditions during real-world tasks offers new possibilities for identifying condition-specific biomarkers at scale.

## Ethics Statement

The Chesapeake IRB approved the HBN protocol. Participants aged 18+ or guardians of minors provided written informed consent for secondary data sharing. Phenotypic data access was governed by a CMI-CUNY Data Transfer and Use Agreement (6/27/2024), and data analysis was approved by CUNY’s IRB under protocol 2024-0867-CCNY (12/14/2024).

## Data Availability

Phenotypical and diagnostic data are available upon request and with the establishment of a Data Use Agreement with the Child Mind Institute at https://fcon_1000.projects.nitrc.org/indi/cmi_healthy_brain_network/.

## Acknowledgments & Funding

NIH-NIMH P50 MH109429, CA Department of Health, grant to Nathan Kline Institute, Milham (PI).

## Notes

### Competing Interest Statement

The authors have declared no competing interest.

