## Supplementary Materials for "Gaze and Pupil Patterns in ADHD and ASD during Naturalistic Viewing"

**Section S2. Alternative Analysis Methods** **7**

**Section S3. Linear mixed-effects model results** **9**

**Bibliography** **16**

**Table S1: Detailed Statistical Test Results for Table 1**

| Variable | ADHD-Inatt vs TD | ADHD-Hyper vs TD | ASD(ADHD+) vs TD | ASD(ADHD−) vs TD | Other vs TD | ASD(ADHD+) vs ASD(ADHD−) |
|---|---|---|---|---|---|---|
| Age | $Z = 3.66$<br>$p = 0.001$ | $Z = -1.37$<br>$p = 0.854$ | $Z = 2.80$<br>$p = 0.026$ | $Z = 2.54$<br>$p = 0.055$ | $Z = 4.15$<br>$p < 0.001$ | $Z = -0.97$<br>$p = 0.334$ |
| Sex | $\chi^2(1) = 27.83$<br>$p < 0.001$ | $\chi^2(1) = 48.75$<br>$p < 0.001$ | $\chi^2(1) = 56.50$<br>$p < 0.001$ | $\chi^2(1) = 23.54$<br>$p < 0.001$ | $\chi^2(1) = 0.03$<br>$p = 1.000$ | $\chi^2(1) = 0.02$<br>$p = 0.899$ |
| Inattention | $Z = 18.15$<br>$p < 0.001$ | $Z = 15.68$<br>$p < 0.001$ | $Z = 13.41$<br>$p < 0.001$ | $Z = 3.66$<br>$p = 0.001$ | $Z = 2.46$<br>$p = 0.069$ | $t(270.0) = 6.80$<br>$p < 0.001$ |
| Hyperactivity | $Z = 14.74$<br>$p < 0.001$ | $Z = 18.94$<br>$p < 0.001$ | $Z = 13.64$<br>$p < 0.001$ | $Z = 3.77$<br>$p < 0.001$ | $Z = 0.47$<br>$p = 1.000$ | $t(270.0) = 6.93$<br>$p < 0.001$ |
| Autism | $Z = 10.95$<br>$p < 0.001$ | $Z = 12.19$<br>$p < 0.001$ | $Z = 15.89$<br>$p < 0.001$ | $Z = 10.67$<br>$p < 0.001$ | $Z = 4.25$<br>$p < 0.001$ | $t(269.0) = 1.34$<br>$p = 0.182$ |
| Full-Scale | $t(1443.0) = -7.84$<br>$p < 0.001$ | $t(922.0) = -5.50$<br>$p < 0.001$ | $t(429.0) = -6.63$<br>$p < 0.001$ | $t(294.0) = -5.95$<br>$p < 0.001$ | $t(806.0) = -5.47$<br>$p < 0.001$ | $t(247.0) = 1.04$<br>$p = 0.298$ |
| Verbal | $Z = -5.88$<br>$p < 0.001$ | $Z = -3.74$<br>$p < 0.001$ | $t(429.0) = -6.22$<br>$p < 0.001$ | $t(294.0) = -6.05$<br>$p < 0.001$ | $t(806.0) = -5.06$<br>$p < 0.001$ | $t(247.0) = 1.61$<br>$p = 0.109$ |
| Validity (%) | $Z = -1.91$<br>$p = 0.281$ | $Z = -2.21$<br>$p = 0.135$ | $Z = -0.87$<br>$p = 1.000$ | $Z = -1.27$<br>$p = 1.000$ | $Z = -2.57$<br>$p = 0.050$ | $Z = 0.75$<br>$p = 0.451$ |

Statistical tests. $t$: Independent $t$-tests (normal continuous variables), $Z$: Wilcoxon rank-sum tests (non-normal continuous variables; via Kolmogorov-Smirnov normality test), and $\chi^2$: Chi-square tests (categorical variables).

**Table S2: Adjusted odds ratios among diagnostic statuses, age and sex.** A. Associations between diagnostic statuses. Each aOR is symmetric: the average of the log odds ratios from the two logistic regressions in which each status predicts the other, each adjusted for the third status, age (per year) and sex. B. aORs for age (per year) and sex from the logistic regression predicting each diagnostic status, adjusted for the other two statuses. Values are aOR [95% CI] ($N = 2421$). ADHD-Combined presentation is coded as both Inattentive and Hyperactive. * $p < 0.01$, ** $p < 0.001$, Bonferroni-corrected across three tests.

**A**

| Pair | aOR [95% CI] | $p$ (Bonferroni) |
|---|---|---|
| ADHD-Inatt – ADHD-Hyper | 10.70 [8.38, 13.66]** | 3.40E-80 |
| ADHD-Inatt – ASD | 1.62 [1.20, 2.17]* | 0.0043 |
| ADHD-Hyper – ASD | 1.22 [0.90, 1.65] | 0.58 |

**B**

| Outcome | N cases | Age (per year) | Female |
|---|---|---|---|
| ADHD-Inatt | 1274 | 1.08 [1.05, 1.11]** | 0.75 [0.62, 0.90]* |
| ADHD-Hyper | 706 | 0.81 [0.79, 0.84]** | 0.53 [0.43, 0.66]** |
| ASD | 280 | 1.06 [1.02, 1.10]* | 0.30 [0.22, 0.42]** |

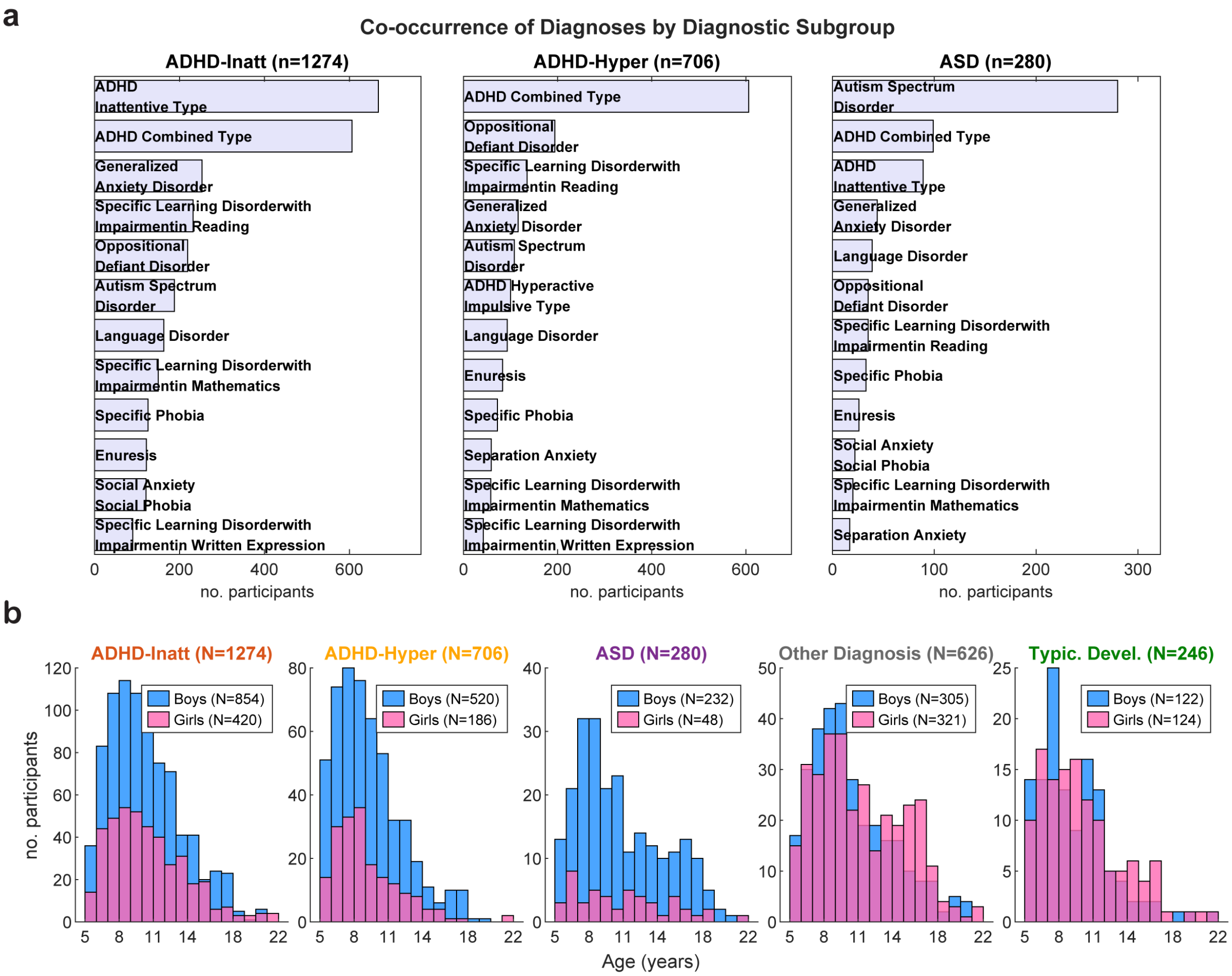

**Figure S1:** **Co-occurring diagnoses and age distributions.** a) Co-occurring diagnoses (Top 12) in each of the three diagnostic groups of interest. b) Age-Sex distribution across the diagnostic groups. ADHD-Inatt and ADHD-Hyper contain participants with an ADHD-Combined presentation. Other Diagnosis group contains participants with some DSM-5 positive diagnosis, excluding those with ADHD or ASD.

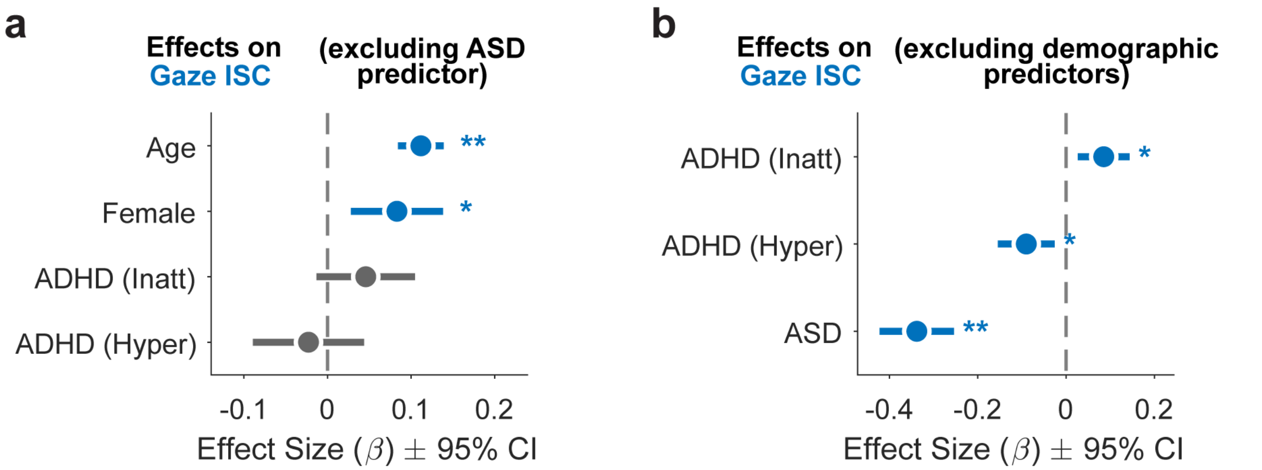

**Figure S2: Unmodeled clinical and demographic covariates produce spurious gaze typicality effects.** Simulated unadjusted designs reveal that evaluating overlapping phenotypes in isolation generates statistical artifacts. (a) Forest plot of standardized multivariate effect sizes ($\beta\pm$ 95% CI) for Gaze-ISC when the co-occurring ASD predictor is excluded from the model. Omitting ASD produces a spurious sex effect (Female, * $p < 0.01$), driven by the lower prevalence of ASD in females, while both ADHD effects remain non-significant. (b) Forest plot of effect sizes when demographic predictors (age and sex) are excluded. Omitting age yields spurious, opposite-signed effects for ADHD presentations: ADHD-Inattentive status artificially predicts more typical gaze (* $p < 0.01$), whereas ADHD-Hyperactive status artificially predicts less typical gaze (* $p < 0.01$). These artifactual effects mirror the underlying developmental skew of the presentations, as hyperactivity declines and gaze typicality increases with age. Significant associations are indicated in blue (* $p < 0.01$, ** $p < 0.001$), and non-significant associations in gray.

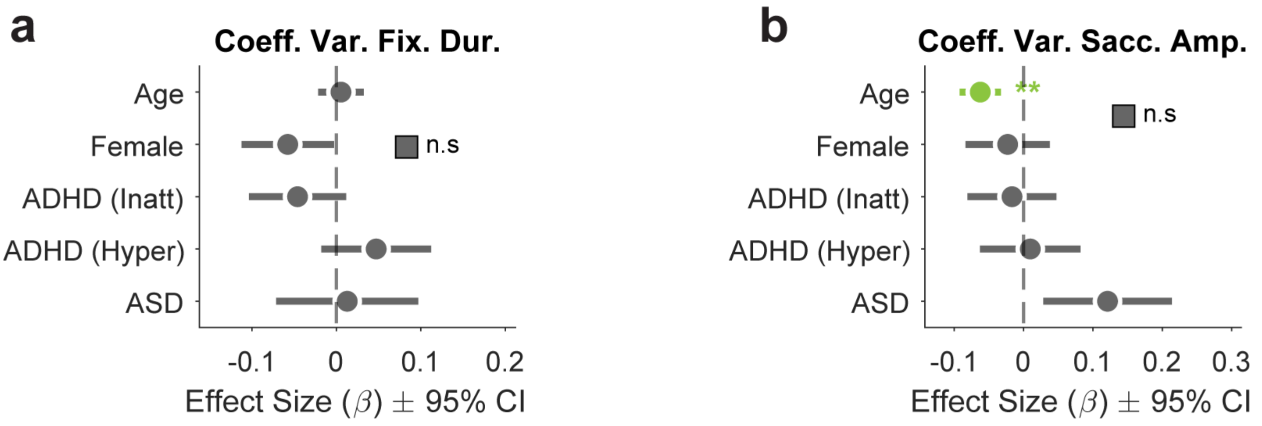

**Figure S3: Within-participant variability of basic oculomotor behaviors is not altered by diagnostic status.** Forest plots detailing the standardized multivariate effect sizes ($\beta\pm$ 95% CI) for the coefficient of variation (CV) of basic oculomotor features. (a) For fixation duration CV, all demographic and clinical predictors, yielded non-significant effects (n.s., Bonferroni corrected for 2 comparisons). (b) For saccadic amplitude CV, older age is significantly associated with reduced variability ($p < 0.001$, highlighted in green, Bonferroni corrected for 2 comparisons). However, no diagnostic dimension or sex showed a significant association with saccadic amplitude variability. All non-significant associations across both panels are depicted in gray. Consistent with the mean-level oculomotor results, these findings confirm that the relative dispersion and intrinsic fluctuation of basic oculomotor control remain typical across both ASD and ADHD.

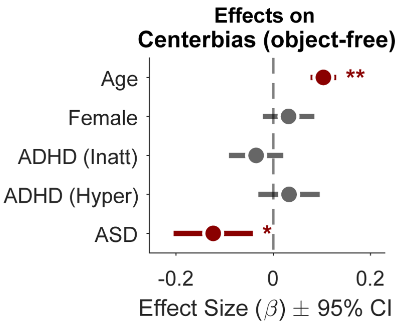

**Figure S4: Reduced center bias in ASD persists during object-free scenes.** Forest plot of standardized multivariate effect sizes ($\beta\pm$ 95% CI) predicting center bias, calculated exclusively during scenes lacking central regions of interest (object-free). Consistent with the full-stimulus analysis, an ASD diagnosis remains significantly associated with reduced center bias (* $p < 0.01$). This confirms that the dispersed spatial viewing strategy in ASD is a fundamental tendency, rather than merely a by-product of looking away from centrally presented social content. Older age remains significantly associated with increased center bias ($p < 0.001$). ADHD-Inattentive status, ADHD-Hyperactive status, and sex (Female) show no significant associations. Significant effects are colored in dark red, while non-significant effects are shown in gray.

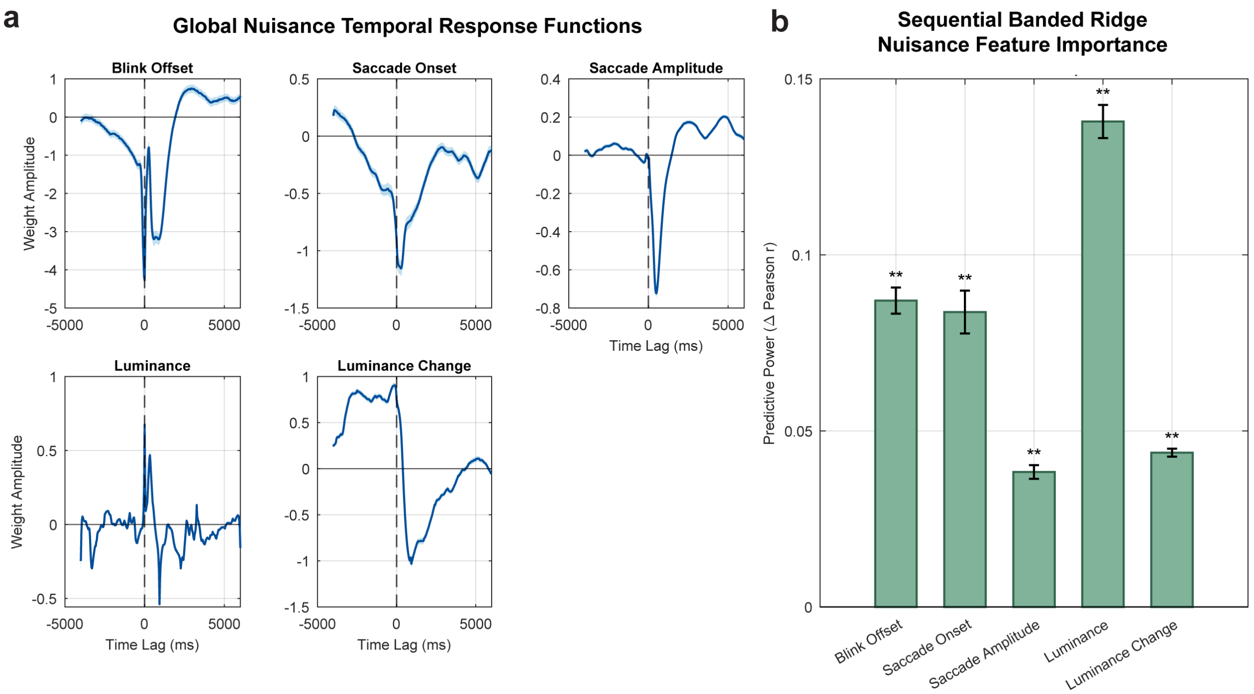

**Figure S5: Nuisance temporal response functions (TRFs) and their contribution to pupil prediction.** (a) Group-mean TRFs (shading, 95% CI across participants) for each nuisance regressor, in the order they entered the model. Weights are in units of z-scored pupil per unit regressor. Blink and saccade responses resemble previous studies[1, 2]. Luminance and its derivative jointly capture the pupillary light reflex. (b) Cross-validated gain in prediction accuracy from each regressor. For each held-out participant, pupil size was predicted from the mean TRFs of all other participants (leave-one-participant-out); accuracy is the Pearson $r$ between predicted and recorded pupil, averaged over that participant's videos. Bars show the increase in $r$ from adding each regressor to the model containing the regressors to its left; the first bar is the accuracy of the blink-offset-only model. Error bars = mean ± 95% CI. **$\Delta r > 0$ in all 1,000 bootstrap samples ($p \leq 0.001$; participants resampled with replacement, and videos resampled within each participant). Acausal components in the TRFs are probably due to interpolation artifacts.

### Section S1. Supplementary Methods

#### S1.1 Participant selection and quality criteria

A total of 4,429 participants took part in the eye-tracking experiment (Releases 1–11), of whom 3,145 had valid eye-tracking files for at least one of the four videos (1,523 for all four) and 2,547 met signal quality criteria (see S1.2). Excluding participants for whom clinicians recommended further testing to rule out ASD left 2,509 participants with clinician-confirmed diagnoses or confirmed typical development. Of these, 2,421 had valid screen resolutions (1440 × 900 or 800 × 600) and formed the final sample. Eye-tracking data were recorded at 30 Hz ($n = 360$), 60 Hz ($n = 1{,}783$), or 120 Hz ($n = 278$).

Stimuli were sourced from: "Despicable Me" (http://www.youtube.com/watch?v=HNXxJIhVALI), "Fun with Fractals" (http://www.youtube.com/watch?v=XwWyTts06tU), "Diary of a Wimpy Kid" (http://www.youtube.com/watch?v=7ZVEIgPeDCE), and "The Present" (https://vimeo.com/152985022).

#### S1.2 Eye-tracking pre-processing

Invalid gaze samples were identified using the eye tracker's native pupil confidence metric, retaining only maximum-confidence samples. Missing samples were reconstructed via linear interpolation, with nearest-neighbor extrapolation at endpoints. Video views were excluded if they contained below 65% valid data pre-interpolation, off-stimulus gaze for over 65% of the duration, or total durations deviating by more than 0.5 s from the per-stimulus median. Valid time series were resampled to 30 Hz.

#### S1.3 Blink, Saccade, and Fixation Detection

Event detection used the eye with higher device-detected validity. Blinks were identified with default eye-tracker settings. Fixations and saccades were detected from linearly interpolated 3D gaze vectors (re-normalized for accurate dispersion calculations) using a moving-window Identification by Dispersion-Threshold (I-DT) algorithm[3], defining fixations as periods where maximum angular dispersion remained at or below 1.6° for at least 75 ms[4]. Only intervals composed of raw, non-interpolated samples were retained as valid fixations, with fixation location computed as the mean 2D screen coordinates across the interval.

Saccades were defined between consecutive valid fixations, with amplitude computed as the total angular displacement between 3D gaze vectors from the end of one fixation to the start of the next. For each stimulus, we computed the mean and coefficient of variation of fixation duration and saccadic amplitude, alongside blink rate derived from the mean inter-blink interval.

#### S1.4 Pupil pre-processing and Temporal Response Functions

Following interpolation, we removed light-reflex and oculomotor artifacts from continuous pupil signals using Temporal Response Function (TRF) modeling[1, 2]. TRFs model the continuous pupil signal $y(t)$ as a sum of convolutions between $F$ predictors $x_f(t)$ and response weights $w_f(\tau)$ across time lags $\tau$:

$$y(t) = \sum_{f=1}^{F} \sum_{\tau=\tau_{\min}}^{\tau_{\max}} w_f(\tau)\, x_f(t-\tau) + \epsilon(t) \tag{1}$$

TRFs were fit per participant using the mTRF-Toolbox[5] over lags of −4 to 6 s to capture full oculomotor[2] and pupillary light-reflex dynamics[6]. Group-mean TRFs are available in Fig. S5a. To account for collinearity among naturalistic predictors and assign shared variance to mechanical artifacts first, we used Sequential Banded Ridge Regression[7, 8, 9], adding nuisance features in fixed order: binary blink offsets, binary saccade onsets, saccade amplitude, continuous log-luminance, and luminance change. For each feature, 15 log-spaced regularization penalties ($\lambda$ from $10^{-1}$ to $10^{5}$) were evaluated via leave-one-subject-out cross-validation to maximize out-of-sample Pearson correlation with the pupil signal (cross-validated prediction gains in Fig. S5b). Each feature was scaled by its optimal $\lambda$ and held constant as subsequent features were added, ensuring that additional variance was uniquely explained by each new predictor. Subtracting the modeled nuisance responses from $y(t)$ yielded the residual $\epsilon(t)$ as the artifact-cleaned pupil time series.

### Section S2. Alternative Analysis Methods

To further validate our methodological decisions, we conducted several additional post-hoc sensitivity analyses:

##### Including IQ

Because typically developing participants had higher IQ than diagnostic groups (Table 1), we refit all models including full-scale IQ as a covariate ($N = 2{,}272$ with available IQ). Higher IQ was associated with more typical gaze (Gaze-ISC $\beta = 0.154$), more looking at social ROIs, and higher Pupil-ISC ($\beta = 0.086$). Diagnostic associations were largely preserved. ASD remained associated with lower Gaze-ISC ($\beta = -0.275$) and reduced looking at characters, faces, and mouths ($\beta = -0.15$ to $-0.18$), with typical oculomotor measures and face latency. Both ASD ($\beta = -0.126$) and ADHD-Hyperactive status ($\beta = -0.080$) remained associated with lower Pupil-ISC, and the lower $1/f$ exponent remained specific to ADHD-Hyperactive status ($\beta = -0.120$). The dissociation in Pupil-ISC was preserved: adding Gaze-ISC attenuated the ASD effect by 52% and the hyperactivity effect by 9%, whereas adding the $1/f$ exponent attenuated the hyperactivity effect by 19% and did not attenuate the ASD effect. One additional effect emerged: ADHD-Inattentive status was associated with slightly higher Gaze-ISC ($\beta = 0.082$, $p = 0.006$), likely reflecting suppression by the lower IQ of inattentive participants. Being opposite in sign to a deficit, it does not alter the conclusion that no ADHD presentation was associated with reduced gaze typicality.

##### Non-additive effects of co-occurring ASD and ADHD

To test whether ASD and ADHD combine non-additively, we added ASD $\times$ ADHD-Inattentive and ASD $\times$ ADHD-Hyperactive interaction terms to the main model. ASD $\times$ ADHD-Hyperactive was not significant for any outcome. ASD $\times$ ADHD-Inattentive was significant for two outcomes. For Gaze-ISC ($\beta = 0.437$), the reduction in gaze typicality was larger in autistic participants without inattentive ADHD ($\beta = -0.610$) than in those with it ($\beta = -0.17$). For the variability of saccade amplitude ($\beta = -0.343$), autistic participants without inattentive ADHD showed more variable saccade amplitudes ($\beta = 0.351$). No other interaction was significant (all $p \geq 0.01$). ADHD main effects remained non-significant for all gaze measures.

##### Mutually exclusive diagnostic groups

We also modeled diagnoses as mutually exclusive groups (ADHD-only, ASD-only, ASD+ADHD), relative to participants with neither diagnosis, with the same covariates. ADHD-only was not associated with any gaze measure (Gaze-ISC $\beta = -0.005$). Both ASD groups showed lower Gaze-ISC (ASD-only $\beta = -0.647$; ASD+ADHD $\beta = -0.212$), reduced looking at characters, faces and mouths ($\beta = -0.18$ to $-0.26$), typical face latency, and lower Pupil-ISC ($\beta = -0.224$ and $-0.142$), which was attenuated by Gaze-ISC ($-71$% and $-37$%) but not by the $1/f$ exponent. More variable saccade amplitudes appeared only in ASD-only ($\beta = 0.352$). Pooling ADHD presentations removed the pupil effects specific to hyperactivity: ADHD-only was not associated with Pupil-ISC ($\beta = -0.037$, $p = 0.085$) or the $1/f$ exponent ($\beta = 0.019$). This shows how combining presentations dilutes an effect carried by hyperactivity.

Together, these analyses support the main conclusions. Atypical gaze is specific to ASD, and the reduction in Pupil-ISC in ASD depends on gaze. They also indicate that the magnitude of gaze atypicality, and the presence of oculomotor variability, differ between autistic participants with and without co-occurring ADHD. Given the small ASD-only group ($n = 68$), these subgroup differences require replication.

##### Removing 'Other Diagnoses' altogether

To test whether including participants with diagnoses other than ASD or ADHD affected our conclusions, we refit all models after excluding them, so that diagnostic effects were estimated relative to typically developing participants only ($N = 1{,}702$ with valid data, of whom 246 typically developing). Participants with ASD or ADHD were retained regardless of other co-occurring diagnoses. All main associations were preserved with similar effect sizes. ASD was associated with lower Gaze-ISC ($\beta = -0.350$), reduced looking at characters, faces and mouths ($\beta = -0.19$ to $-0.23$), and lower Pupil-ISC ($\beta = -0.146$), with typical oculomotor measures, orienting latency, and face latency. Neither ADHD presentation was associated with any gaze measure. ADHD-Hyperactive status remained associated with lower Pupil-ISC ($\beta = -0.070$) and a lower $1/f$ exponent ($\beta = -0.162$). The dissociation in Pupil-ISC was preserved: adding Gaze-ISC attenuated the ASD effect by 57% but the hyperactivity effect by only 1%, whereas adding the $1/f$ exponent attenuated the hyperactivity effect by 25% and did not attenuate the ASD effect. Our conclusions therefore do not depend on the composition of the non-ASD, non-ADHD reference group.

##### Age × Diagnosis Interactions

Our main mixed-effects model estimates each diagnostic effect as a constant difference across ages. Such an average difference can be detected even if it changes with age. To test whether diagnostic effects varied with age, we added interactions between age (mean-centered) and each diagnostic status to the full model. No interaction reached statistical significance for any outcome (all $p > 0.01$). The closest were Age × ASD for center bias ($p = 0.010$) and for Pupil-ISC ($p = 0.021$). Diagnostic effects estimated at the mean age were virtually unchanged: ASD on Gaze-ISC $\beta = -0.341$; ASD on Pupil-ISC $\beta = -0.148$; ADHD-Hyperactive on Pupil-ISC $\beta = -0.073$; ADHD-Hyperactive on the $1/f$ exponent $\beta = -0.152$. The dissociation in Pupil-ISC was preserved. Within the age range studied (5–22 years), we therefore found no evidence that the diagnostic effects reported here depend on age, although the cross-sectional design limits sensitivity to developmental change.

##### Continuous trait measures

We replaced the three diagnostic statuses with continuous parent-rated traits: SWAN inattention, SWAN hyperactivity, and ASSQ autistic traits ($N = 2{,}292$). Results were similar with the SRS-2 in place of the ASSQ ($N = 2{,}005$). Effects on the attention side were reproduced. Higher autistic traits were associated with lower Gaze-ISC ($\beta = -0.084$ per SD), reduced looking at characters, faces, and mouths ($\beta = -0.07$ to $-0.08$), reduced center bias ($\beta = -0.058$), and lower Pupil-ISC ($\beta = -0.038$). They were not associated with oculomotor measures or with latency after scene cuts. Higher autistic traits were associated with slightly slower orienting to faces ($\beta = 0.039$, $p = 0.0099$); this effect was positive but non-significant in all diagnosis-based models. ADHD traits were not associated with reduced gaze typicality. In contrast to diagnostic status, hyperactivity traits were associated with neither Pupil-ISC ($\beta = -0.015$, $p = 0.27$) nor the $1/f$ exponent ($\beta = -0.025$, $p = 0.24$). The pupil differences we report may therefore reflect clinically significant hyperactivity, as captured by a clinician's diagnosis, rather than variation along the full range of parent-rated hyperactive traits.

### Section S3. Linear mixed-effects model results

##### Results for Feature: Gaze-ISC

Formula: `ISC_gaze ~ 1 + Age + Female + positive_inattentive_ADHD + positive_hyperactive_ADHD + positive_autism + fs + stim + (1 | subject_id)`
Observations: $N = 6535$ | Unique Subjects: 2421

| Predictor | Estimate ($\beta$) | SE | t-stat | DF | 95% CI | p-value |
|---|---|---|---|---|---|---|
| (Intercept) | -0.001 | 0.0414 | -0.02 | 6524 | [-0.0821, 0.0801] | 0.98 |
| Age | 0.1177 | 0.0138 | 8.54 | 6524 | [0.0907, 0.1447] | 1.59E-17 |
| Female | 0.048 | 0.0281 | 1.71 | 6524 | [-0.007, 0.103] | 0.0874 |
| ADHD-Inatt | 0.0603 | 0.0296 | 2.03 | 6524 | [0.0022, 0.1184] | 0.0419 |
| ADHD-Hyper | -0.015 | 0.0335 | -0.45 | 6524 | [-0.0806, 0.0506] | 0.653 |
| ASD | -0.344 | 0.0428 | -8.03 | 6524 | [-0.428, -0.26] | 1.17E-15 |
| fs_60 | -0.0324 | 0.0371 | -0.87 | 6524 | [-0.1053, 0.0404] | 0.382 |
| fs_120 | -0.3212 | 0.0564 | -5.69 | 6524 | [-0.4318, -0.2106] | 1.31E-08 |
| stim_FF | -1.0739 | 0.0282 | -38.11 | 6524 | [-1.1292, -1.0187] | 3.81E-287 |
| stim_TP | 0.7058 | 0.0242 | 29.12 | 6524 | [0.6583, 0.7533] | 2.53E-175 |
| stim_WK | 0.0493 | 0.0216 | 2.28 | 6524 | [0.007, 0.0916] | 0.0224 |

##### Results for Feature: # Fixations

Formula: `no_fixations ~ 1 + Age + Female + positive_inattentive_ADHD + positive_hyperactive_ADHD + positive_autism + fs + stim + (1 | subject_id)`
Observations: $N = 6535$ | Unique Subjects: 2421

| Predictor | Estimate ($\beta$) | SE | t-stat | DF | 95% CI | p-value | p-value (Bonferroni) |
|---|---|---|---|---|---|---|---|
| (Intercept) | -0.6797 | 0.0421 | -16.16 | 6524 | [-0.7622, -0.5973] | 1.22E-57 | 4.87E-57 |
| Age | 0.0334 | 0.0143 | 2.34 | 6524 | [0.0054, 0.0615] | 0.0195 | 0.0779 |
| Female | 0.0164 | 0.0293 | 0.56 | 6524 | [-0.0411, 0.0739] | 0.576 | 1 |
| ADHD-Inatt | 0.0212 | 0.0311 | 0.68 | 6524 | [-0.0398, 0.0822] | 0.495 | 1 |
| ADHD-Hyper | 0.0112 | 0.035 | 0.32 | 6524 | [-0.0575, 0.0798] | 0.75 | 1 |
| ASD | 0.0279 | 0.0445 | 0.63 | 6524 | [-0.0593, 0.1152] | 0.531 | 1 |
| fs_60 | 0.8523 | 0.0393 | 21.68 | 6524 | [0.7752, 0.9294] | 1.11E-100 | 4.46E-100 |
| fs_120 | 1.3034 | 0.0562 | 23.18 | 6524 | [1.1932, 1.4136] | 2.57E-114 | 1.03E-113 |
| stim_FF | 0.0468 | 0.0153 | 3.06 | 6524 | [0.0168, 0.0767] | 0.00221 | 0.00884 |
| stim_TP | 0.6986 | 0.0131 | 53.44 | 6524 | [0.6729, 0.7242] | 0 | 0 |
| stim_WK | -0.8401 | 0.0115 | -72.81 | 6524 | [-0.8627, -0.8175] | 0 | 0 |

##### Results for Feature: Mean Fix. Dur.

Formula: `mean_fix_duration ~ 1 + Age + Female + positive_inattentive_ADHD + positive_hyperactive_ADHD + positive_autism + fs + stim + (1 | subject_id)`
Observations: $N = 6535$ | Unique Subjects: 2421

| Predictor | Estimate ($\beta$) | SE | t-stat | DF | 95% CI | p-value | p-value (Bonferroni) |
|---|---|---|---|---|---|---|---|
| (Intercept) | 1.0979 | 0.0506 | 21.69 | 6524 | [0.9986, 1.1971] | 9.29E-101 | 3.71E-100 |
| Age | -0.0348 | 0.0173 | -2.02 | 6524 | [-0.0687, -0.001] | 0.0435 | 0.174 |
| Female | -0.0763 | 0.0354 | -2.15 | 6524 | [-0.1457, -0.0069] | 0.0312 | 0.125 |
| ADHD-Inatt | -0.0053 | 0.0376 | -0.14 | 6524 | [-0.079, 0.0684] | 0.889 | 1 |
| ADHD-Hyper | -0.0433 | 0.0423 | -1.02 | 6524 | [-0.1262, 0.0396] | 0.306 | 1 |
| ASD | -0.0023 | 0.0537 | -0.04 | 6524 | [-0.1076, 0.1029] | 0.965 | 1 |
| fs_60 | -0.9702 | 0.0476 | -20.4 | 6524 | [-1.0635, -0.877] | 1.00E-89 | 4.00E-89 |
| fs_120 | -1.932 | 0.0675 | -28.63 | 6524 | [-2.0643, -1.7997] | 6.44E-170 | 2.58E-169 |
| stim_FF | -0.3854 | 0.0152 | -25.43 | 6524 | [-0.4152, -0.3557] | 4.33E-136 | 1.73E-135 |
| stim_TP | -0.1237 | 0.013 | -9.54 | 6524 | [-0.1491, -0.0983] | 1.90E-21 | 7.58E-21 |
| stim_WK | -0.2242 | 0.0114 | -19.61 | 6524 | [-0.2466, -0.2018] | 3.02E-83 | 1.21E-82 |

##### Results for Feature: Mean Sacc. Amp.

Formula: `mean_sacc_amplitude ~ 1 + Age + Female + positive_inattentive_ADHD + positive_hyperactive_ADHD + positive_autism + fs + stim + (1 | subject_id)`
Observations: $N = 6535$ | Unique Subjects: 2421

| Predictor | Estimate ($\beta$) | SE | t-stat | DF | 95% CI | p-value | p-value (Bonferroni) |
|---|---|---|---|---|---|---|---|
| (Intercept) | 1.0635 | 0.0495 | 21.49 | 6524 | [0.9665, 1.1605] | 4.49E-99 | 1.80E-98 |
| Age | -0.088 | 0.0168 | -5.24 | 6524 | [-0.1209, -0.055] | 1.66E-07 | 6.65E-07 |
| Female | -0.0245 | 0.0344 | -0.71 | 6524 | [-0.092, 0.0429] | 0.475 | 1 |
| ADHD-Inatt | -0.0124 | 0.0365 | -0.34 | 6524 | [-0.0839, 0.0591] | 0.734 | 1 |
| ADHD-Hyper | -0.0238 | 0.0411 | -0.58 | 6524 | [-0.1043, 0.0567] | 0.562 | 1 |
| ASD | 0.0009 | 0.0522 | 0.02 | 6524 | [-0.1015, 0.1033] | 0.987 | 1 |
| fs_60 | -1.4372 | 0.046 | -31.23 | 6524 | [-1.5274, -1.347] | 1.05E-199 | 4.19E-199 |
| fs_120 | -1.1731 | 0.0663 | -17.68 | 6524 | [-1.3031, -1.043] | 2.24E-68 | 8.95E-68 |
| stim_FF | 0.2623 | 0.0208 | 12.61 | 6524 | [0.2215, 0.3031] | 4.89E-36 | 1.96E-35 |
| stim_TP | 0.033 | 0.0178 | 1.85 | 6524 | [-0.0019, 0.0679] | 0.0639 | 0.256 |
| stim_WK | 0.3178 | 0.0157 | 20.2 | 6524 | [0.2869, 0.3486] | 4.70E-88 | 1.88E-87 |

##### Results for Feature: Blink Rate

Formula: `mean_blink_rate_per_min ~ 1 + Age + Female + positive_inattentive_ADHD + positive_hyperactive_ADHD + positive_autism + fs + stim + (1 | subject_id)`
Observations: $N = 6457$ | Unique Subjects: 2406

| Predictor | Estimate ($\beta$) | SE | t-stat | DF | 95% CI | p-value | p-value (Bonferroni) |
|---|---|---|---|---|---|---|---|
| (Intercept) | 0.0764 | 0.0581 | 1.31 | 6446 | [-0.0375, 0.1903] | 0.189 | 0.755 |
| Age | 0.1718 | 0.0197 | 8.74 | 6446 | [0.1333, 0.2104] | 3.04E-18 | 1.21E-17 |
| Female | -0.0695 | 0.0404 | -1.72 | 6446 | [-0.1486, 0.0097] | 0.0854 | 0.342 |
| ADHD-Inatt | -0.1014 | 0.0428 | -2.37 | 6446 | [-0.1853, -0.0175] | 0.0179 | 0.0715 |
| ADHD-Hyper | 0.0752 | 0.0482 | 1.56 | 6446 | [-0.0194, 0.1697] | 0.119 | 0.476 |
| ASD | -0.0689 | 0.0614 | -1.12 | 6446 | [-0.1894, 0.0515] | 0.262 | 1 |
| fs_60 | 0.0321 | 0.054 | 0.59 | 6446 | [-0.0738, 0.138] | 0.552 | 1 |
| fs_120 | -0.151 | 0.0782 | -1.93 | 6446 | [-0.3044, 0.0023] | 0.0536 | 0.214 |
| stim_FF | 0.5841 | 0.0243 | 23.99 | 6446 | [0.5364, 0.6318] | 7.10E-122 | 2.84E-121 |
| stim_TP | -0.1249 | 0.0209 | -5.99 | 6446 | [-0.1658, -0.084] | 2.26E-09 | 9.05E-09 |
| stim_WK | -0.1693 | 0.0185 | -9.15 | 6446 | [-0.2056, -0.133] | 7.67E-20 | 3.07E-19 |

##### Results for Feature: Coeff. Var. Sacc. Amp.

Formula: `cv_sacc_amplitude ~ 1 + Age + Female + positive_inattentive_ADHD + positive_hyperactive_ADHD + positive_autism + fs + stim + (1 | subject_id)`
Observations: $N = 6535$ | Unique Subjects: 2421

| Predictor | Estimate ($\beta$) | SE | t-stat | DF | 95% CI | p-value | p-value (Bonferroni) |
|---|---|---|---|---|---|---|---|
| (Intercept) | -0.6471 | 0.0457 | -14.15 | 6524 | [-0.7368, -0.5575] | 8.64E-45 | 1.73E-44 |
| Age | -0.0623 | 0.0152 | -4.1 | 6524 | [-0.0922, -0.0325] | 4.18E-05 | 8.36E-05 |
| Female | -0.0229 | 0.031 | -0.74 | 6524 | [-0.0836, 0.0379] | 0.46 | 0.921 |
| ADHD-Inatt | -0.0166 | 0.0327 | -0.51 | 6524 | [-0.0808, 0.0475] | 0.611 | 1 |
| ADHD-Hyper | 0.0098 | 0.0369 | 0.26 | 6524 | [-0.0626, 0.0822] | 0.792 | 1 |
| ASD | 0.1212 | 0.0473 | 2.56 | 6524 | [0.0284, 0.2139] | 0.0105 | 0.021 |
| fs_60 | 0.702 | 0.041 | 17.12 | 6524 | [0.6216, 0.7824] | 2.58E-64 | 5.15E-64 |
| fs_120 | 1.3436 | 0.0624 | 21.53 | 6524 | [1.2212, 1.4659] | 2.02E-99 | 4.04E-99 |
| stim_FF | 0.4961 | 0.0315 | 15.77 | 6524 | [0.4345, 0.5578] | 4.90E-55 | 9.80E-55 |
| stim_TP | -0.0775 | 0.0271 | -2.86 | 6524 | [-0.1306, -0.0244] | 0.0042 | 0.00841 |
| stim_WK | -0.1075 | 0.0241 | -4.46 | 6524 | [-0.1547, -0.0602] | 8.42E-06 | 1.68E-05 |

##### Results for Feature: Coeff. Var. Fix. Dur.

Formula: `cv_fix_duration ~ 1 + Age + Female + positive_inattentive_ADHD + positive_hyperactive_ADHD + positive_autism + fs + stim + (1 | subject_id)`
Observations: $N = 6535$ | Unique Subjects: 2421

| Predictor | Estimate ($\beta$) | SE | t-stat | DF | 95% CI | p-value | p-value (Bonferroni) |
|---|---|---|---|---|---|---|---|
| (Intercept) | 0.3736 | 0.0427 | 8.74 | 6524 | [0.2898, 0.4573] | 2.86E-18 | 5.72E-18 |
| Age | 0.0055 | 0.0138 | 0.4 | 6524 | [-0.0215, 0.0325] | 0.691 | 1 |
| Female | -0.0575 | 0.0279 | -2.06 | 6524 | [-0.1122, -0.0028] | 0.0393 | 0.0785 |
| ADHD-Inatt | -0.0459 | 0.0293 | -1.57 | 6524 | [-0.1035, 0.0116] | 0.118 | 0.235 |
| ADHD-Hyper | 0.0471 | 0.0332 | 1.42 | 6524 | [-0.0179, 0.1122] | 0.155 | 0.311 |
| ASD | 0.0129 | 0.0428 | 0.3 | 6524 | [-0.0711, 0.0969] | 0.764 | 1 |
| fs_60 | -0.2787 | 0.0366 | -7.61 | 6524 | [-0.3505, -0.2069] | 3.17E-14 | 6.35E-14 |
| fs_120 | -0.5216 | 0.059 | -8.84 | 6524 | [-0.6373, -0.4059] | 1.23E-18 | 2.47E-18 |
| stim_FF | -0.0259 | 0.0381 | -0.68 | 6524 | [-0.1005, 0.0488] | 0.497 | 0.994 |
| stim_TP | -0.1701 | 0.0331 | -5.15 | 6524 | [-0.235, -0.1053] | 2.74E-07 | 5.49E-07 |
| stim_WK | -0.1334 | 0.0298 | -4.48 | 6524 | [-0.1918, -0.0751] | 7.47E-06 | 1.49E-05 |

##### Results for Feature: Fixation Latency ($\log_{10}$ ms)

Formula: `logFixLat ~ 1 + Age + Female + positive_inattentive_ADHD + positive_hyperactive_ADHD + positive_autism + fs + stim + (1 | subject_id)`
Observations: $N = 6535$ | Unique Subjects: 2421

| Predictor | Estimate ($\beta$) | SE | t-stat | DF | 95% CI | p-value |
|---|---|---|---|---|---|---|
| (Intercept) | 0.4265 | 0.0443 | 9.63 | 6524 | [0.3397, 0.5134] | 8.46E-22 |
| Age | -0.0403 | 0.0145 | -2.78 | 6524 | [-0.0687, -0.0118] | 0.00551 |
| Female | -0.0345 | 0.0295 | -1.17 | 6524 | [-0.0923, 0.0232] | 0.241 |
| ADHD-Inatt | -0.0149 | 0.031 | -0.48 | 6524 | [-0.0758, 0.0459] | 0.631 |
| ADHD-Hyper | -0.0021 | 0.0351 | -0.06 | 6524 | [-0.0708, 0.0666] | 0.952 |
| ASD | -0.0482 | 0.0451 | -1.07 | 6524 | [-0.1367, 0.0402] | 0.285 |
| fs_60 | -0.5312 | 0.0388 | -13.7 | 6524 | [-0.6072, -0.4552] | 4.04E-42 |
| fs_120 | -0.8662 | 0.0609 | -14.22 | 6524 | [-0.9855, -0.7468] | 3.15E-45 |
| stim_FF | -0.0172 | 0.0357 | -0.48 | 6524 | [-0.0872, 0.0527] | 0.629 |
| stim_TP | 0.0555 | 0.0308 | 1.8 | 6524 | [-0.005, 0.1159] | 0.0721 |
| stim_WK | 0.1902 | 0.0276 | 6.89 | 6524 | [0.1361, 0.2444] | 6.29E-12 |

**Results for Feature: Centerbias**

Formula: `cb_prop5 ~ 1 + Age + Female + positive_inattentive_ADHD + positive_hyperactive_ADHD + positive_autism + fs + stim + (1 | subject_id)`

Observations: $N = 6535$ | Unique Subjects: 2421

| Predictor | Estimate ($\beta$) | SE | t-stat | DF | 95% CI | p-value |
|---|---|---|---|---|---|---|
| (Intercept) | 0.1813 | 0.0413 | 4.39 | 13055 | [0.1003, 0.2622] | 1.14E-05 |
| Age | 0.0728 | 0.0141 | 5.15 | 6524 | [0.0451, 0.1005] | 2.72E-07 |
| Female | 0.0243 | 0.0287 | 0.85 | 6524 | [-0.032, 0.0805] | 0.398 |
| ADHD-Inatt | -0.0327 | 0.0302 | -1.08 | 6524 | [-0.0919, 0.0266] | 0.28 |
| ADHD-Hyper | 0.0476 | 0.0342 | 1.39 | 6524 | [-0.0194, 0.1145] | 0.164 |
| ASD | -0.1282 | 0.044 | -2.91 | 6524 | [-0.2145, -0.042] | 0.00358 |
| fs_60 | 0.1001 | 0.0378 | 2.65 | 6524 | [0.0261, 0.1741] | 0.00801 |
| fs_120 | -0.4081 | 0.0598 | -6.83 | 6524 | [-0.5253, -0.291] | 9.27E-12 |
| stim_FF | -0.3163 | 0.0361 | -8.75 | 6524 | [-0.3871, -0.2454] | 2.66E-18 |
| stim_TP | -0.0912 | 0.0313 | -2.91 | 6524 | [-0.1525, -0.0299] | 0.00357 |
| stim_WK | 0.381 | 0.0281 | 13.58 | 6524 | [0.326, 0.436] | 1.99E-41 |

**Results for Feature: Pixel-level (Itti-Koch)**

Formula: `salPixel ~ 1 + Age + Female + positive_inattentive_ADHD + positive_hyperactive_ADHD + positive_autism + fs + stim + (1 | subject_id)`

Observations: $N = 6534$ | Unique Subjects: 2421

| Predictor | Estimate ($\beta$) | SE | t-stat | DF | 95% CI | p-value | p-value (Bonferroni) |
|---|---|---|---|---|---|---|---|
| (Intercept) | -0.5574 | 0.0303 | -18.38 | 6523 | [-0.6169, -0.498] | 1.43E-73 | 1.43E-73 |
| Age | 0.0103 | 0.0096 | 1.07 | 6523 | [-0.0085, 0.0292] | 0.282 | 0.282 |
| Female | 0.0211 | 0.0195 | 1.09 | 6523 | [-0.017, 0.0593] | 0.278 | 0.278 |
| ADHD-Inatt | -0.0211 | 0.0204 | -1.03 | 6523 | [-0.0612, 0.019] | 0.303 | 0.303 |
| ADHD-Hyper | 0.0055 | 0.0231 | 0.24 | 6523 | [-0.0398, 0.0508] | 0.812 | 0.812 |
| ASD | -0.047 | 0.03 | -1.57 | 6523 | [-0.1057, 0.0118] | 0.117 | 0.117 |
| fs_60 | 0.0041 | 0.0255 | 0.16 | 6523 | [-0.046, 0.0541] | 0.873 | 0.873 |
| fs_120 | -0.1168 | 0.042 | -2.78 | 6523 | [-0.1991, -0.0344] | 0.00545 | 0.00545 |
| stim_FF | -0.4773 | 0.029 | -16.45 | 6523 | [-0.5342, -0.4204] | 1.31E-59 | 1.31E-59 |
| stim_TP | 0.9736 | 0.0253 | 38.47 | 6523 | [0.924, 1.0233] | 5.60E-292 | 5.60E-292 |
| stim_WK | 1.2763 | 0.0229 | 55.79 | 6523 | [1.2314, 1.3211] | 0 | 0 |

**Results for Feature: Characters**

Formula: `pref_Characters ~ 1 + Age + Female + positive_inattentive_ADHD + positive_hyperactive_ADHD + positive_autism + fs + stim + (1 | subject_id)`

Observations: $N = 5579$ | Unique Subjects: 2389

| Predictor | Estimate ($\beta$) | SE | t-stat | DF | 95% CI | p-value | p-value (Bonferroni) |
|---|---|---|---|---|---|---|---|
| (Intercept) | -0.0083 | 0.0445 | -0.19 | 5569 | [-0.0954, 0.0789] | 0.852 | 1 |
| Age | 0.0613 | 0.0148 | 4.15 | 5569 | [0.0323, 0.0902] | 3.34E-05 | 0.0002 |
| Female | 0.0011 | 0.03 | 0.04 | 5569 | [-0.0577, 0.0599] | 0.972 | 1 |
| ADHD-Inatt | -0.0033 | 0.0317 | -0.11 | 5569 | [-0.0655, 0.0588] | 0.916 | 1 |
| ADHD-Hyper | 0.019 | 0.0357 | 0.53 | 5569 | [-0.0511, 0.089] | 0.596 | 1 |
| ASD | -0.2102 | 0.0459 | -4.58 | 5569 | [-0.3002, -0.1201] | 4.86E-06 | 2.91E-05 |
| fs_60 | 0.0632 | 0.0392 | 1.61 | 5569 | [-0.0137, 0.14] | 0.107 | 0.642 |
| fs_120 | -0.4709 | 0.0624 | -7.55 | 5569 | [-0.5932, -0.3485] | 5.23E-14 | 3.14E-13 |
| stim_TP | -0.4821 | 0.0299 | -16.14 | 5569 | [-0.5407, -0.4235] | 2.63E-57 | 1.58E-56 |
| stim_WK | 0.3378 | 0.0268 | 12.63 | 5569 | [0.2853, 0.3902] | 4.75E-36 | 2.85E-35 |

##### Results for Feature: Faces

Formula: `pref_Faces ~ 1 + Age + Female + positive_inattentive_ADHD + positive_hyperactive_ADHD + positive_autism + fs + stim + (1 | subject_id)`
Observations: $N = 5579$ | Unique Subjects: 2389

| Predictor | Estimate ($\beta$) | SE | t-stat | DF | 95% CI | p-value | p-value (Bonferroni) |
|---|---|---|---|---|---|---|---|
| (Intercept) | 0.2972 | 0.0457 | 6.51 | 5569 | [0.2077, 0.3868] | 8.32E-11 | 4.99E-10 |
| Age | 0.0667 | 0.0152 | 4.38 | 5569 | [0.0369, 0.0965] | 1.20E-05 | 7.19E-05 |
| Female | 0.0023 | 0.0309 | 0.08 | 5569 | [-0.0583, 0.063] | 0.94 | 1 |
| ADHD-Inatt | -0.0153 | 0.0327 | -0.47 | 5569 | [-0.0795, 0.0488] | 0.639 | 1 |
| ADHD-Hyper | 0.0202 | 0.0369 | 0.55 | 5569 | [-0.0521, 0.0925] | 0.584 | 1 |
| ASD | -0.2121 | 0.0474 | -4.48 | 5569 | [-0.305, -0.1192] | 7.75E-06 | 4.65E-05 |
| fs_60 | -0.0536 | 0.0406 | -1.32 | 5569 | [-0.1331, 0.0259] | 0.186 | 1 |
| fs_120 | -0.5495 | 0.0639 | -8.6 | 5569 | [-0.6748, -0.4242] | 1.07E-17 | 6.40E-17 |
| stim_TP | -0.7188 | 0.0294 | -24.47 | 5569 | [-0.7764, -0.6612] | 1.07E-125 | 6.40E-125 |
| stim_WK | -0.0389 | 0.0263 | -1.48 | 5569 | [-0.0904, 0.0125] | 0.138 | 0.829 |

##### Results for Feature: Mouth

Formula: `pref_Mouth ~ 1 + Age + Female + positive_inattentive_ADHD + positive_hyperactive_ADHD + positive_autism + fs + stim + (1 | subject_id)`
Observations: $N = 5579$ | Unique Subjects: 2389

| Predictor | Estimate ($\beta$) | SE | t-stat | DF | 95% CI | p-value | p-value (Bonferroni) |
|---|---|---|---|---|---|---|---|
| (Intercept) | 0.3038 | 0.0409 | 7.43 | 5569 | [0.2236, 0.3839] | 1.25E-13 | 7.52E-13 |
| Age | 0.0735 | 0.0137 | 5.36 | 5569 | [0.0466, 0.1003] | 8.74E-08 | 5.24E-07 |
| Female | -0.016 | 0.0279 | -0.57 | 5569 | [-0.0707, 0.0388] | 0.567 | 1 |
| ADHD-Inatt | -0.012 | 0.0296 | -0.4 | 5569 | [-0.07, 0.046] | 0.686 | 1 |
| ADHD-Hyper | 0.0031 | 0.0333 | 0.09 | 5569 | [-0.0622, 0.0684] | 0.925 | 1 |
| ASD | -0.1884 | 0.0427 | -4.41 | 5569 | [-0.2721, -0.1047] | 1.05E-05 | 6.30E-05 |
| fs_60 | -0.1163 | 0.0368 | -3.16 | 5569 | [-0.1884, -0.0441] | 0.00159 | 0.00954 |
| fs_120 | -0.4796 | 0.0568 | -8.44 | 5569 | [-0.591, -0.3682] | 3.95E-17 | 2.37E-16 |
| stim_TP | -1.1148 | 0.0236 | -47.23 | 5569 | [-1.1611, -1.0686] | 0 | 0 |
| stim_WK | 0.3453 | 0.021 | 16.42 | 5569 | [0.3041, 0.3865] | 3.13E-59 | 1.88E-58 |

##### Results for Feature: Eyes

Formula: `pref_Eyes ~ 1 + Age + Female + positive_inattentive_ADHD + positive_hyperactive_ADHD + positive_autism + fs + stim + (1 | subject_id)`
Observations: $N = 5578$ | Unique Subjects: 2389

| Predictor | Estimate ($\beta$) | SE | t-stat | DF | 95% CI | p-value | p-value (Bonferroni) |
|---|---|---|---|---|---|---|---|
| (Intercept) | 0.0839 | 0.0502 | 1.67 | 5568 | [-0.0144, 0.1822] | 0.0944 | 0.566 |
| Age | -0.0007 | 0.0168 | -0.04 | 5568 | [-0.0337, 0.0322] | 0.965 | 1 |
| Female | 0.0303 | 0.0343 | 0.88 | 5568 | [-0.0369, 0.0975] | 0.377 | 1 |
| ADHD-Inatt | -0.0354 | 0.0363 | -0.98 | 5568 | [-0.1066, 0.0357] | 0.329 | 1 |
| ADHD-Hyper | 0.0482 | 0.0409 | 1.18 | 5568 | [-0.032, 0.1284] | 0.238 | 1 |
| ASD | -0.0767 | 0.0524 | -1.46 | 5568 | [-0.1795, 0.026] | 0.143 | 0.859 |
| fs_60 | -0.044 | 0.0452 | -0.97 | 5568 | [-0.1326, 0.0445] | 0.33 | 1 |
| fs_120 | -0.3203 | 0.0697 | -4.59 | 5568 | [-0.457, -0.1836] | 4.47E-06 | 2.68E-05 |
| stim_TP | -0.2065 | 0.0289 | -7.14 | 5568 | [-0.2632, -0.1498] | 1.08E-12 | 6.50E-12 |
| stim_WK | 0.0492 | 0.0258 | 1.91 | 5568 | [-0.0014, 0.0997] | 0.0565 | 0.339 |

##### Results for Feature: Text

Formula: `pref_Text ~ 1 + Age + Female + positive_inattentive_ADHD + positive_hyperactive_ADHD + positive_autism + fs + stim + (1 | subject_id)`
Observations: $N = 3007$ | Unique Subjects: 2224

| Predictor | Estimate ($\beta$) | SE | t-stat | DF | 95% CI | p-value | p-value (Bonferroni) |
|---|---|---|---|---|---|---|---|
| (Intercept) | 0.9957 | 0.0489 | 20.36 | 2998 | [0.8999, 1.0916] | 1.91E-86 | 1.15E-85 |
| Age | 0.1288 | 0.0151 | 8.56 | 2998 | [0.0993, 0.1583] | 1.81E-17 | 1.09E-16 |
| Female | 0.0086 | 0.0304 | 0.28 | 2998 | [-0.051, 0.0681] | 0.778 | 1 |
| ADHD-Inatt | 0.0171 | 0.0319 | 0.54 | 2998 | [-0.0455, 0.0797] | 0.592 | 1 |
| ADHD-Hyper | 0.0258 | 0.0361 | 0.71 | 2998 | [-0.0451, 0.0966] | 0.476 | 1 |
| ASD | -0.0543 | 0.0468 | -1.16 | 2998 | [-0.1461, 0.0375] | 0.246 | 1 |
| fs_60 | -0.0209 | 0.041 | -0.51 | 2998 | [-0.1013, 0.0596] | 0.611 | 1 |
| fs_120 | -0.112 | 0.0649 | -1.73 | 2998 | [-0.2392, 0.0153] | 0.0846 | 0.507 |
| stim_WK | -1.3859 | 0.0298 | -46.52 | 2998 | [-1.4443, -1.3275] | 0 | 0 |

##### Results for Feature: Background

Formula: `pref_Background ~ 1 + Age + Female + positive_inattentive_ADHD + positive_hyperactive_ADHD + positive_autism + fs + stim + (1 | subject_id)`
Observations: $N = 6530$ | Unique Subjects: 2420

| Predictor | Estimate ($\beta$) | SE | t-stat | DF | 95% CI | p-value | p-value (Bonferroni) |
|---|---|---|---|---|---|---|---|
| (Intercept) | -0.4695 | 0.011 | -42.84 | 6519 | [-0.491, -0.448] | 0 | 0 |
| Age | -0.0113 | 0.0035 | -3.18 | 6519 | [-0.0182, -0.0043] | 0.00149 | 0.00896 |
| Female | -0.0056 | 0.0072 | -0.78 | 6519 | [-0.0197, 0.0085] | 0.434 | 1 |
| ADHD-Inatt | 0.0043 | 0.0076 | 0.57 | 6519 | [-0.0105, 0.0192] | 0.566 | 1 |
| ADHD-Hyper | -0.0018 | 0.0086 | -0.21 | 6519 | [-0.0186, 0.015] | 0.833 | 1 |
| ASD | 0.0434 | 0.011 | 3.94 | 6519 | [0.0218, 0.065] | 8.40E-05 | 0.000504 |
| fs_60 | -0.0014 | 0.0095 | -0.15 | 6519 | [-0.0199, 0.0172] | 0.884 | 1 |
| fs_120 | 0.0969 | 0.0151 | 6.41 | 6519 | [0.0673, 0.1265] | 1.55E-10 | 9.30E-10 |
| stim_FF | 2.7976 | 0.0095 | 294.21 | 6519 | [2.779, 2.8163] | 0 | 0 |
| stim_TP | 0.2751 | 0.0082 | 33.36 | 6519 | [0.259, 0.2913] | 2.02E-225 | 1.21E-224 |
| stim_WK | -0.0221 | 0.0074 | -2.99 | 6519 | [-0.0367, -0.0076] | 0.00283 | 0.017 |

##### Results for Feature: Face Latency ($\log_{10}$ ms)

Formula: `logFaceLat ~ 1 + Age + Female + positive_inattentive_ADHD + positive_hyperactive_ADHD + positive_autism + fs + stim + (1 | subject_id)`
Observations: $N = 5583$ | Unique Subjects: 2391

| Predictor | Estimate ($\beta$) | SE | t-stat | DF | 95% CI | p-value |
|---|---|---|---|---|---|---|
| (Intercept) | 0.4467 | 0.0436 | 10.24 | 5573 | [0.3611, 0.5322] | 2.19E-24 |
| Age | -0.0299 | 0.0144 | -2.07 | 5573 | [-0.0581, -0.0016] | 0.0381 |
| Female | -0.0132 | 0.0292 | -0.45 | 5573 | [-0.0705, 0.0441] | 0.651 |
| ADHD-Inatt | -0.0296 | 0.0309 | -0.96 | 5573 | [-0.0901, 0.0309] | 0.338 |
| ADHD-Hyper | -0.0598 | 0.0348 | -1.72 | 5573 | [-0.128, 0.0085] | 0.0861 |
| ASD | 0.037 | 0.0448 | 0.83 | 5573 | [-0.0509, 0.1249] | 0.409 |
| fs_60 | -0.0799 | 0.038 | -2.1 | 5573 | [-0.1544, -0.0053] | 0.0357 |
| fs_120 | 0.3711 | 0.0615 | 6.03 | 5573 | [0.2505, 0.4918] | 1.73E-09 |
| stim_TP | -0.455 | 0.0313 | -14.54 | 5573 | [-0.5164, -0.3937] | 4.55E-47 |
| stim_WK | -0.6648 | 0.0281 | -23.66 | 5573 | [-0.7199, -0.6097] | 5.66E-118 |

##### Results for Feature: Pupil-ISC

Formula: `ISC_cleanedpupil ~ 1 + Age + Female + positive_inattentive_ADHD + positive_hyperactive_ADHD + positive_autism + fs + stim + (1 | subject_id)`

Observations: $N = 6535$ | Unique Subjects: 2421

| Predictor | Estimate ($\beta$) | SE | t-stat | DF | 95% CI | p-value |
|---|---|---|---|---|---|---|
| (Intercept) | -0.8634 | 0.0316 | -27.35 | 6524 | [-0.9252, -0.8015] | 5.02E-156 |
| Age | -0.0518 | 0.0104 | -4.97 | 6524 | [-0.0722, -0.0314] | 6.76E-07 |
| Female | -0.0377 | 0.0212 | -1.78 | 6524 | [-0.0792, 0.0038] | 0.0751 |
| ADHD-Inatt | 0.0215 | 0.0223 | 0.96 | 6524 | [-0.0223, 0.0653] | 0.336 |
| ADHD-Hyper | -0.0755 | 0.0252 | -2.99 | 6524 | [-0.1249, -0.0261] | 0.00276 |
| ASD | -0.1409 | 0.0324 | -4.35 | 6524 | [-0.2044, -0.0774] | 1.38E-05 |
| fs_60 | -0.0197 | 0.0279 | -0.71 | 6524 | [-0.0745, 0.035] | 0.48 |
| fs_120 | -0.1901 | 0.0433 | -4.39 | 6524 | [-0.2749, -0.1053] | 1.13E-05 |
| stim_FF | 0.3002 | 0.0239 | 12.58 | 6524 | [0.2535, 0.347] | 7.25E-36 |
| stim_TP | 1.4166 | 0.0206 | 68.8 | 6524 | [1.3762, 1.4569] | 0 |
| stim_WK | 1.6885 | 0.0184 | 91.78 | 6524 | [1.6524, 1.7246] | 0 |

##### Results for Feature: Mean pupil

Formula: `mean_pupil ~ 1 + Age + Female + positive_inattentive_ADHD + positive_hyperactive_ADHD + positive_autism + fs + stim + (1 | subject_id)`

Observations: $N = 6535$ | Unique Subjects: 2421

| Predictor | Estimate ($\beta$) | SE | t-stat | DF | 95% CI | p-value |
|---|---|---|---|---|---|---|
| (Intercept) | 0.596 | 0.0561 | 10.62 | 6524 | [0.4859, 0.706] | 4.14E-26 |
| Age | -0.1581 | 0.0192 | -8.25 | 6524 | [-0.1957, -0.1205] | 1.93E-16 |
| Female | 0.0354 | 0.0394 | 0.9 | 6524 | [-0.0417, 0.1126] | 0.368 |
| ADHD-Inatt | -0.0167 | 0.0418 | -0.4 | 6524 | [-0.0987, 0.0652] | 0.689 |
| ADHD-Hyper | -0.0525 | 0.047 | -1.12 | 6524 | [-0.1447, 0.0396] | 0.264 |
| ASD | 0.0608 | 0.0597 | 1.02 | 6524 | [-0.0561, 0.1777] | 0.308 |
| fs_60 | -0.42 | 0.053 | -7.93 | 6524 | [-0.5238, -0.3162] | 2.52E-15 |
| fs_120 | -0.8825 | 0.0747 | -11.82 | 6524 | [-1.0288, -0.7361] | 6.61E-32 |
| stim_FF | -0.5911 | 0.0133 | -44.47 | 6524 | [-0.6172, -0.5651] | 0 |
| stim_TP | -0.2047 | 0.0114 | -18.01 | 6524 | [-0.227, -0.1824] | 7.88E-71 |
| stim_WK | -0.213 | 0.01 | -21.27 | 6524 | [-0.2327, -0.1934] | 4.15E-97 |

##### Results for Feature: $1/f$ Exponent ($\chi$)

Formula: `exponent ~ 1 + Age + Female + positive_inattentive_ADHD + positive_hyperactive_ADHD + positive_autism + fs + stim + (1 | subject_id)`

Observations: $N = 6535$ | Unique Subjects: 2421

| Predictor | Estimate ($\beta$) | SE | t-stat | DF | 95% CI | p-value |
|---|---|---|---|---|---|---|
| (Intercept) | -0.6034 | 0.0479 | -12.59 | 6524 | [-0.6974, -0.5095] | 6.31E-36 |
| Age | -0.0676 | 0.0162 | -4.17 | 6524 | [-0.0994, -0.0358] | 3.07E-05 |
| Female | 0.0503 | 0.0332 | 1.51 | 6524 | [-0.0148, 0.1153] | 0.13 |
| ADHD-Inatt | 0.0865 | 0.0352 | 2.46 | 6524 | [0.0176, 0.1555] | 0.0139 |
| ADHD-Hyper | -0.1497 | 0.0396 | -3.78 | 6524 | [-0.2273, -0.072] | 0.000159 |
| ASD | 0.0656 | 0.0505 | 1.3 | 6524 | [-0.0333, 0.1645] | 0.194 |
| fs_60 | 0.2931 | 0.0443 | 6.62 | 6524 | [0.2063, 0.38] | 3.99E-11 |
| fs_120 | -0.4387 | 0.0645 | -6.8 | 6524 | [-0.5651, -0.3123] | 1.11E-11 |
| stim_FF | -0.3137 | 0.0228 | -13.74 | 6524 | [-0.3585, -0.269] | 2.33E-42 |
| stim_TP | 0.52 | 0.0196 | 26.59 | 6524 | [0.4817, 0.5584] | 5.80E-148 |
| stim_WK | 0.9009 | 0.0173 | 52.06 | 6524 | [0.8669, 0.9348] | 0 |
